# Nuclear Mitochondrial Sequences in Great Ape Telomere-to-Telomere Genomes

**DOI:** 10.1101/2025.04.24.650511

**Authors:** Edmundo Torres-González, Marzia A. Cremona, Jessica M. Storer, Mario Ventura, Rachel J. O’Neill, Kateryna D. Makova

**Affiliations:** Department of Biology, The Pennsylvania State University, PA, USA; Huck Life Sciences Institutes, The Pennsylvania State University, PA, USA; Department of Operations and Decision Systems, Université Laval, Québec, Canada; CHU de Québec-Université Laval Research Center, Québec, Canada; Department of Molecular and Cellular Biology, University of Connecticut, CT, USA; University of Bari Aldo Moro, Bari, Italy; Center for Medical Genomics, The Pennsylvania State University, PA, USA

## Abstract

Mitochondrial sequences have integrated into the nuclear genome since the origin of eukaryotes. Recent insertions that retain homology to extant mitochondrial DNA (mtDNA), termed NUMTs, confound mtDNA sequence analysis. Here, we used great ape Telomere-to-Telomere (T2T) genomes to study NUMTs in bonobo, chimpanzee, human, gorilla, and Bornean and Sumatran orangutans. A phylogeny based on shared and lineage-specific NUMTs accurately recapitulated the great ape species tree topology. NUMTs were enriched at non-functional non-repetitive regions of the nuclear genome, and depleted within enhancers and coding sequences, suggesting negative selection. We validated the presence of a 76-kilobase-long heterozygous NUMT in chimpanzee, which was larger than any other NUMT observed in great apes, and found that dozens of NUMTs on the *Pan* Y chromosome expanded together with palindromes. Our study highlights NUMTs as a dynamic evolutionary force contributing to shaping ape genomes, and is valuable for characterizing mtDNA in great apes.

## Introduction

The nuclear genome has been bombarded with sequences from mitochondria presumably since the origin of the eukaryotic cell. Over evolutionary time, some mitochondrial DNA (mtDNA) fragments integrated into the nuclear genome. As the nucleus depended on mitochondria for energy production and other metabolic pathways, the ability to encode mitochondrial proteins in the nucleus permitted tight regulation of mitochondrial processes. A large fraction of genes in mtDNA moved to the nucleus, an ongoing process that started around 1,200 million years ago (MYA) ^1^. More recent mtDNA integrations in the nuclear genome form nuclear sequences of mitochondrial origin, or NUMTs (i.e., “new-mights”, ^2^), which have a high sequence identity to mtDNA ^3^. Over time, mitochondria became dependent on the nucleus to contribute genes essential to oxidative phosphorylation, replication, and transcription ^4^. Currently, mtDNA harbors just over 30 genes in mammals and most other animals ^5,6^.

Depending on their integration time, NUMTs can be shared by closely related species and contribute to genetic variation between and within them ^7^. NUMTs are ubiquitous and have been previously annotated in human ^8–10^, macaque ^11^, chimpanzee ^12,13^, gorilla, bonobo ^14^, as well as other mammals ^15^, plants, fungi, and protists (reviewed in ^16^). Consistent with a high tolerance of excess sequence in plants, their NUMTs are abundant and can reach hundreds of kilobases (kb) in length ^17^. In contrast, NUMTs in honeybees are relatively short (≤863 base pairs, or bp) as a result of fragmented mtDNA integration ^18^. Human NUMT insertions range from <100 bp ^18^ to the size of a full mtDNA copy (∼16.5 kb in human and other apes) ^19^ and can have a high sequence identity to mtDNA (>99%). NUMTs have been found to integrate non-randomly into the nuclear genome of mammals, preferring transposon-rich and intergenic regions ^15^. Additionally, certain parts of mtDNA are more prone to integration into the nuclear genome as NUMTs ^20^. Note that most of the previous studies analyzed unfinished genome assemblies and thus provided incomplete NUMT annotations.

To insert into the nuclear genome, NUMTs rely on faulty repair mechanisms (e.g., non-homologous end joining) at the sites of double-strand breaks ^21^. Increased frequency of double-strand breaks at the nuclear or mitochondrial genome, e.g., because of induced stress, facilitates NUMT integration ^22^. Recent work in insects suggests that some NUMTs may also integrate into the nuclear genome following reverse transcription ^23^. NUMTs may expand into multiple copies, but they do not encode the machinery to do so alone. This limitation may be overcome by utilizing the mobility of neighboring transposable elements (TEs). TEs are DNA sequences that have the potential to move within the host genome. TEs can be separated into two major classes based on their mobilization intermediate, via replicative (“copy and paste”) or non-replicative transposition (“cut and paste”) ^24^. Long interspersed nuclear elements (LINEs) encode a reverse transcriptase to transpose in the genome, which is also used by their non-autonomous counterparts, short interspersed nuclear elements (SINEs) ^25^. NUMTs could leverage the LINE-encoded reverse transcriptase in a similar way to SINEs. Non-replicative TEs, or DNA transposons, may carry NUMTs when they move across the genome. Therefore, both TEs and NUMTs are expected to be enriched at regions of genomic instability. Alternatively, NUMTS might be passengers of other common large-scale genome rearrangements and expansions ^26^. However, the exact mechanisms of NUMT insertion and expansion have not been fully elucidated.

In some eukaryotes, NUMTs influence nuclear replication and transcription by acting as *cis*-regulatory elements, as observed in *Toxoplasma* with its high rates of NUMT insertions ^27^ and in yeast ^28^. However, NUMT insertions more often disrupt functional regions. In particular, pericentromeric and telomeric regions are prone to NUMT insertions, leading to interrupted centromere activity, aneuploidy, and cellular senescence ^29,30^. NUMT accumulation in brain tissue ^31^ and cancerous cells ^32,33^ have also been linked to early mortality in humans, possibly a remnant of DNA stress.

NUMTs confound mtDNA sequence analysis, whereby NUMT reads align to the mtDNA assembly and are observed as erroneous variation in the mtDNA genome ^34^. Re-sequencing experiments that rely on short reads are especially susceptible to this issue ^35–37^. The improved contiguity of nuclear assemblies generated with long reads helps resolve longer NUMTs and provides a more reliable catalog of NUMTs ^38^.

Recently, the Telomere-to-Telomere (T2T) genome assemblies for human and other great apes were released ^38–41^. This provided us with a unique opportunity to obtain a complete view of NUMTs across the great apes—bonobo (*Pan paniscus*), chimpanzee (*Pan troglodytes*), human (*Homo sapiens*), gorilla (*Gorilla gorilla*), Bornean orangutan (*Pongo pygmaeus*), and Sumatran orangutan (*Pongo abelii*)—with different divergence times. The divergence time is estimated to be ∼2.5 MY between bonobo and chimpanzee (*Pan* genus), ∼7 MY between human and *Pan*, ∼9 MY between gorilla and *Hominini* (human, bonobo, and chimpanzee), ∼1 MY between Bornean and Sumatran orangutans (*Pongo* genus), and ∼17 MYA between *Pongo* and the other great apes, *Homininae* (human, bonobo, and chimpanzee, and gorilla; see ^41^ and references therein).

Here, we used these T2T assemblies to study the integration and expansion dynamics of NUMTs in great ape genomes. We annotated NUMTs in T2T ape genomes and conducted their comparative analysis. We detailed the distribution of NUMTs across the T2T genome for each species and generated a great ape phylogeny based on shared and lineage-specific NUMTs. Then, we tested the contribution of flanking sequence composition to NUMT insertion preferences. Finally, we highlighted examples of NUMTs with intriguing mechanisms of expansion. Our results provide a valuable resource for future studies utilizing ape T2T assemblies, such as the potential functional impact of NUMTs on gene regulation and genome evolution in humans and great apes.

## Results

### NUMTs in ape T2T genomes

We annotated NUMTs in six great ape assemblies with available T2T genomes ^38^. NUMTs were identified separately for each species by a pairwise alignment of mtDNA to the nuclear genome (see Methods and Supplemental Data 1). We provide a detailed description of NUMTs in the primary haploid assembly, and analyze alternate haplotypes when possible (not available for the human CHM13 genome). The NUMTs we detected were at least 30 bp (see Methods) and up to 16.5 kb (a full-length mitochondrial genome) long, except for one 76-kb NUMT on the chimpanzee chromosome homologous to human chromosome 6 (HSA6; here and below we use human chromosome numbering for both human and other species based on homology, i.e., HSA, unless specified otherwise). We detected 713 NUMTs in bonobo, 717 in chimpanzee, 711 in human, 648 in gorilla, 714 in Bornean orangutan, and 715 in Sumatran orangutan for the primary haplotype. Similar numbers of NUMTs were detected for the alternate haplotype (702, 723, 0, 656, 721, and 729, respectively; the human reference genome is haploid and thus does not have the alternate haplotype; Table S1).

We found that NUMTs varied greatly in length within each species but overall were skewed towards smaller events across apes (Figure 1A), with median lengths ranging from 265 to 299 bp. Many NUMTs maintained a high sequence identity to mtDNA (>99%), suggesting recent insertions. Most NUMTs were short (<100 bp), suggesting a bias for short NUMT insertions, potentially confounded by our preferential ability to detect shorter insertions (Figure S1). Short NUMTs with low identity to mtDNA could also be fragments of long, older NUMTs. Analyzing the primary haplotype for each chromosome for each species, we observed a particularly high number of NUMTs on chr2b for *Hominini*, chr17 for *Pongo*, chr22 for human, and chrY for *Pan* (Figure 1B). We hypothesize these high numbers were due to lineage-specific expansions or incomplete lineage sorting, contributing to variation across species.

**Figure 1.**
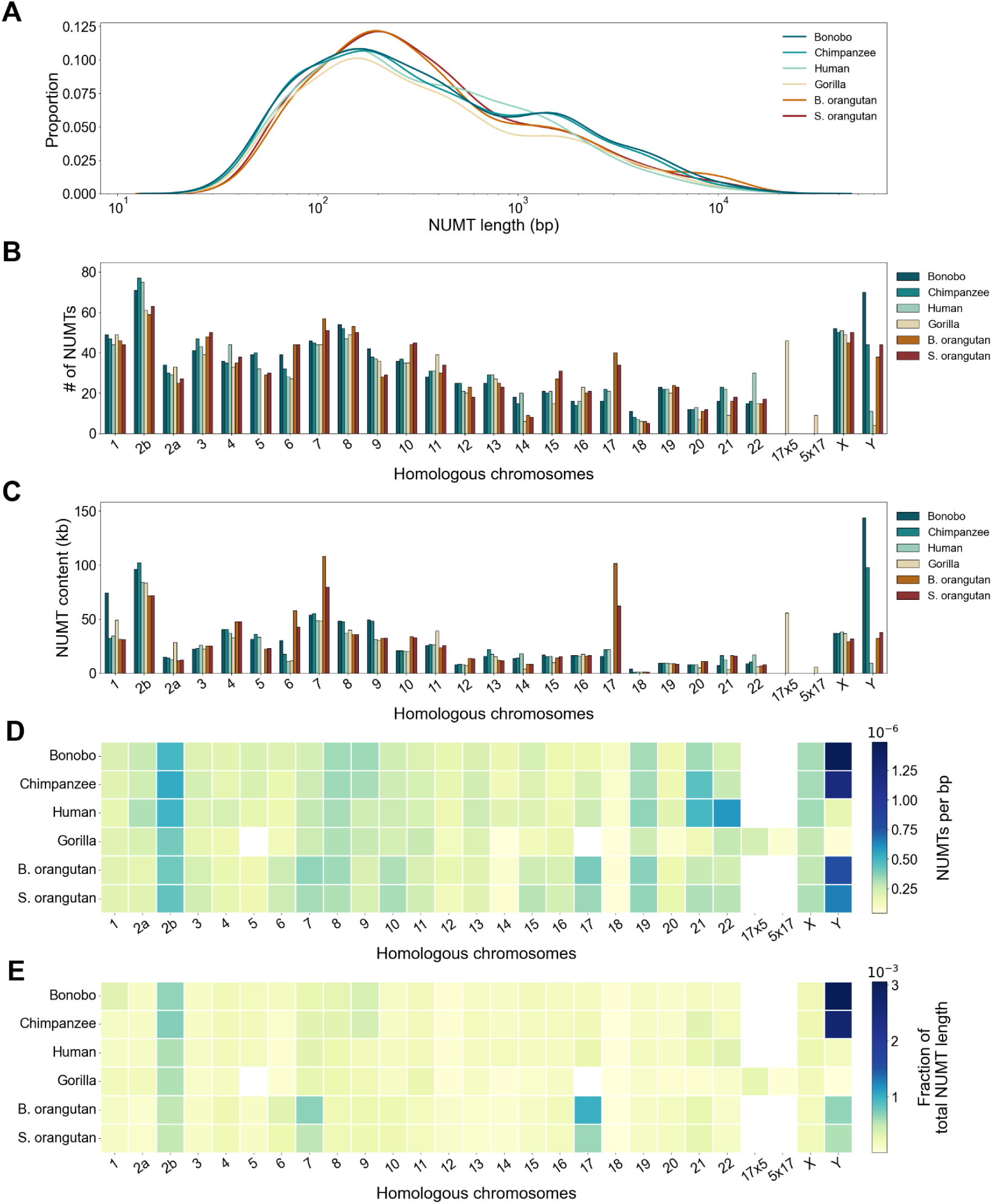
NUMT content across homologous great ape chromosomes in primary haplotype assemblies. Chromosomes were labeled based on their human homolog numbers (HSA). **(A)** Length distribution of NUMT lengths per species. **(B)** The number of NUMTs per chromosome per species. **(C)** NUMT kilobases per chromosome per species (*NUMT content*). **(D)** Number of NUMTs per bp for each chromosome and species. **(E)** The fraction of bases annotated as NUMTs among all bases for each chromosome and species. Human chromosome 2 was divided at the site of an ancient fusion (∼114 megabases, Mb), and each arm was assigned to their homologs HSA2a and HSA2b in great apes. The gorilla chromosomes homologous to HSA5 and HSA17 exchanged genetic material and resulted in two combinations referred to as chr17×5 and chr5×17. They were considered separately from chr5 and chr17 in this analysis.

The total base pairs encompassed by NUMTs (*NUMT content*; Figure 1C) supported the trends observed for NUMT counts (Figure 1B). For instance, chrY in *Pan* had a ∼3-fold higher NUMT density and content than most other chromosomes. Chr1 for bonobo, chr7 and 17 for *Pongo*, and chrY for *Pan* showed a 20- to 100-kb higher NUMT content than homologous chromosomes in other species (Figure 1C).

The alternate assembly showed similar trends to those observed for the primary assembly (Figure S2). However, when considering both haplotypes for *Pongo* (Figures 1 and S1), we noted that differences in NUMT content among orangutan species on chr7 and chr17 were due to inter-haplotype variation (Figures 1 and S2). In addition, the alternate haplotype for chr6 in chimpanzee had a notably higher NUMT content (Figure S2) compared to the primary haplotype (Figure 1C), explained by a single 76-kb NUMT in the alternate haplotype.

### Shared and lineage-specific NUMTs

To identify shared NUMTs across great apes, we aligned NUMT flanks for each species to all genome assemblies using BLAT ^42^. Then we compared the *k*-mer distributions of the query NUMT and the BLAT match, following a method similar to that described in Yoo et al. 2025 ^38^ (see Methods). Natural clades—those that match the species tree topology—were observed most commonly, with 1,300 NUMTs shared by all great apes, 610 between the two *Pongo* species, 508 among *Homininae*, 186 among *Hominini*, and 258 between the two *Pan* species (Figure 2A). Species-specific NUMTs were common in human (N=153) and gorilla (N=118), followed by bonobo (N=65) and chimpanzee (N=62), with the lowest numbers in the orangutan species (N=43 in Sumatran orangutan and N=28 in Bornean orangutan), consistent with their recent divergence from each other. We also observed many clades with a suspected loss of NUMTs in a single species. For instance, many NUMTs were shared across great apes but were absent in either human (N=297) or gorilla (N=191), suggesting deletions. There were also some clades that require at least two events (i.e., multiple NUMT-containing sequence deletions) to explain them (e.g., NUMTs present in African great apes but in neither human nor orangutans), or may be due to incomplete lineage sorting (Figure S3).

**Figure 2.**
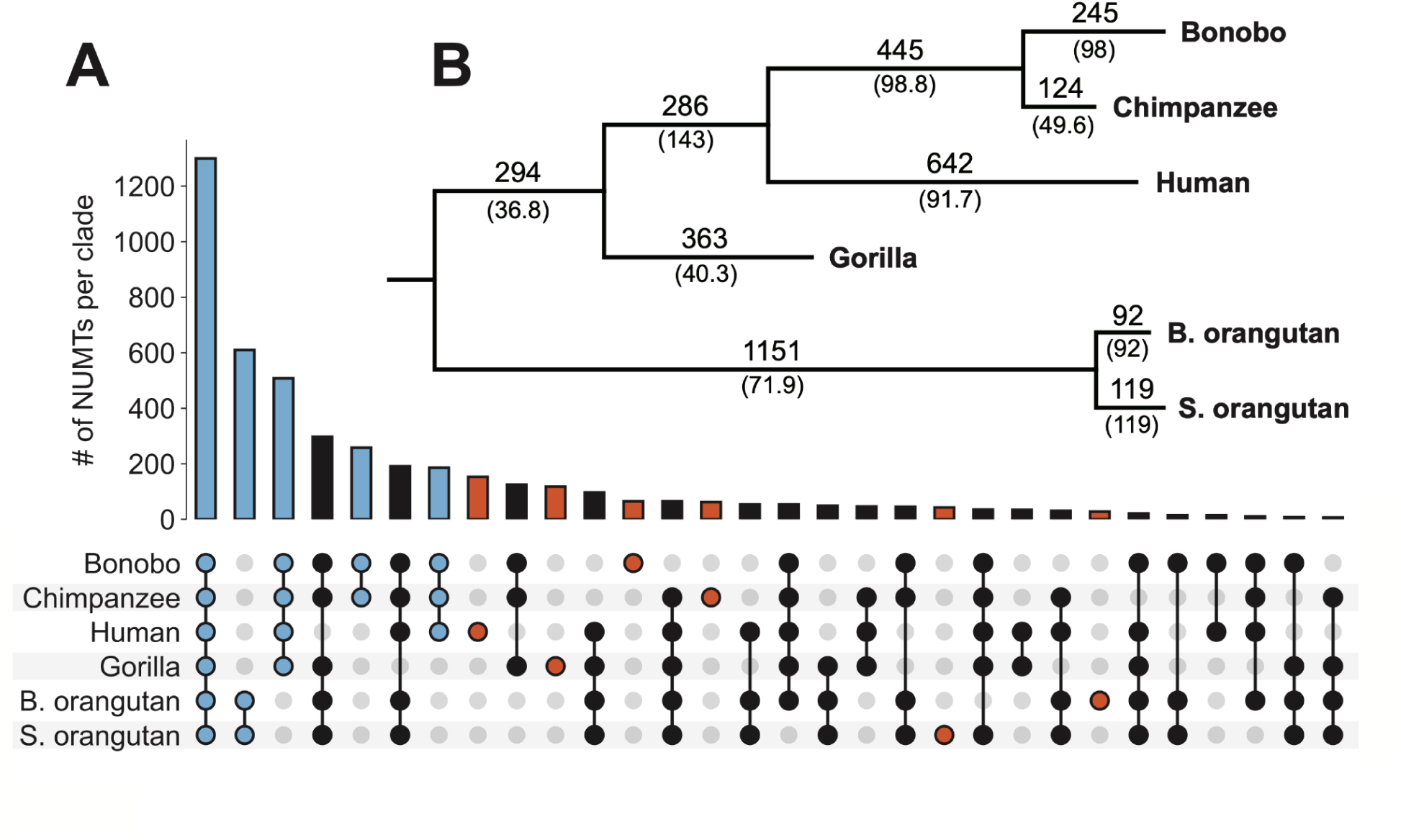
Shared and lineage-specific NUMTs in the primary haplotype. **(A)** Upset plot of shared and lineage-specific NUMT insertions (see text and Methods for details). The 30 most abundant clades were plotted, with species-specific NUMTs in red and those forming natural clades in blue. See Figure S3 for the complete set of clades. **(B)** Maximum parsimony tree of shared and lineage-specific NUMTs. Branches were annotated with the branch length (number of NUMTs that differ between species). The rate of NUMT accumulation, NUMTs per MY, is given in parentheses. A detailed list of which NUMTs are shared was included as Supplemental Data 2.

### Phylogenetic analysis

Using maximum parsimony, we built a phylogenetic tree based on NUMT presence/absence at orthologous locations (Figure 2B). This tree matched the topology of the expected species tree ^38,41^. We computed the rate of NUMT accumulation per species using the tree branch lengths divided by divergence time (divergence times were from Makova et al. 2024 ^41^). Gorilla had the lowest rate of NUMT accumulation (40.3 NUMTs per million years (MY)), followed by chimpanzee (49.6 NUMTs per MY), with higher accumulation rates in human (91.7 NUMTs per MY), bonobo (98 NUMTs per MY), and *Pongo* (92 and 119 NUMTs per MY; Figure 2B). Overall, clades that excluded gorilla displayed higher accumulation rates. For example, the *Hominini* accumulation rate was higher than that of *Homininae*.

### NUMTs across the genome

To determine whether NUMTs were preferentially present in certain genome regions, we studied NUMT presence among functional, repetitive, and presumably neutrally evolving regions (Figure 3). We used the immediate 50-bp flanks upstream and downstream of NUMTs to determine the region in which they integrated.

**Figure 3.**
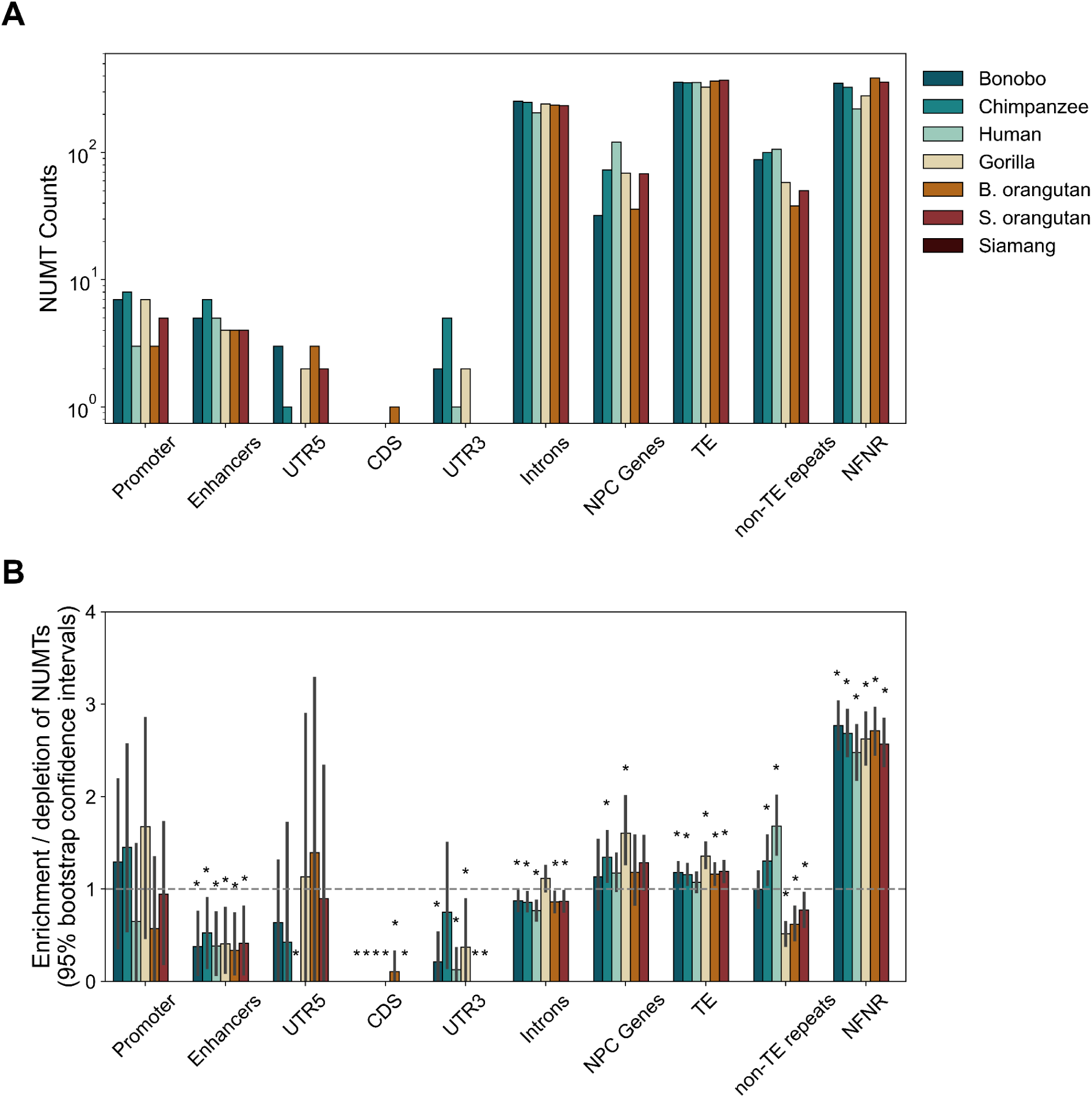
NUMTs at functional, repetitive, and presumably neutrally evolving genome regions. **(A)** The number of NUMTs and **(B)** enrichment in NUMT counts, flanked (50 bp up- or downstream) by each genomic feature. Enrichment scores were computed as the fraction of total NUMTs within the genomic feature divided by the fraction of the genome encompassed by the genomic feature. The horizontal line at 1 represents the genome-wide average. Values above 1 indicate enrichment of NUMTs in that feature, while values below 1 indicate depletion of NUMTs. Significant enrichment or depletion was determined by 95% confidence intervals (obtained with bootstrap, 1,000 repetitions) that do not overlap 1, detailed by a star above the bar. NPC, non-protein-coding genes. NFNR, non-functional non-repetitive regions.

#### Protein-coding genes

NUMTs in coding sequences (CDS) and untranslated regions (UTRs) were infrequent across great apes (Figure 3A). Notably, NUMTs were significantly depleted at 3’UTRs in all species except chimpanzee (bootstrap 95% confidence intervals <1; Figure 3B). CDS regions lacked NUMTs, with the exception of Bornean orangutan (Figure 3B). The observed depletion indicates that NUMTs are not tolerated in coding sequences, suggesting negative selection (Figure 3B). A substantial fraction of all NUMTs, ∼25%, were located within introns across all great apes, comprising ∼120-170 kb in each species (Figure 3A). NUMTs showed a modest, yet significant, 1.3-fold depletion within the introns of all species except gorilla (Figure 3B).

#### Non-protein-coding (NPC) genes and regulatory regions

Non-protein-coding (NPC) genes also housed NUMTs. For instance, NPC genes were significantly enriched in NUMTs for chimpanzee and gorilla (bootstrap 95% confidence intervals >1; Figure 3B). Promoters had NUMT frequencies as expected by chance alone. In contrast, enhancer regions were significantly depleted of NUMTs in all species (Figure 3B), suggesting negative selection.

#### Transposable elements (TEs) and NUMTs

Many NUMTs, 40-60% depending on the species, were within 100 bp of a TE (Figure S4), corresponding to the high fraction of the genome composed of TEs ^38,43^. We observed a modest yet significant NUMT enrichment at TEs in most species (Figure 3B), except in humans.

#### Non-functional non-repetitive regions

A third of all NUMTs were found in non-functional non-repetitive (NFNR) regions of the genome (Figure 3A), defined by excluding all functional ^44^ and RepeatMasker annotations ^45^ for each species. These regions are expected to evolve neutrally. NUMTs for all species were significantly enriched at NFNRs, 2.5- to 2.7-fold in all species (Figure 3B), which may reflect negative selection against NUMTs at functional regions.

### Integration preferences of NUMTs

We used a functional hypothesis testing approach to study potential associations between NUMTs and their sequence context (IWTomics; ^46^). Briefly, we generated a test set of non-overlapping 100-kb regions flanking NUMTs (50-kb upstream and 50-kb downstream), and a control set of 100-kb regions in the rest of the genome. The sequences were annotated in 1-kb increments for functional regions (e.g., origins of replication, see Methods), non-B DNA, CpG islands, and repeat families. We explored differences between NUMTs and control sets for each annotation in human (Figure S5). We observed a depletion of simple repeats at and around NUMTs and a depletion of satellite sequences around NUMTs, which reflects a lack of NUMTs at telomeres or centromeric satellite arrays or a limitation in identifying them in such regions.

### Co-localization of NUMTs and TEs

NUMTs depend on external mechanisms to integrate and expand within the nuclear genome. TE sites are prone to double-strand breaks ^47^, and thus NUMTs may insert in or near TEs. We hypothesized that both TEs and NUMTs are more likely to insert in regions of high genome instability, and thus should co-occur. To test whether NUMTs and TEs co-localize at a large genomic scale, we computed NUMT and TE densities along the genome using two different window sizes (0.1 Mb and 1 Mb; Figure S6). We found that there is a modest negative correlation between TE density and the log of NUMT density in most comparisons (significant in 9 out of 11 comparisons, one comparison showed a significant positive correlation, Pearson’s correlation test; Figure S6), not supporting our hypothesis.

### A large, recent, and heterozygous NUMT insertion in chimpanzee

We identified a 76-kb NUMT array in the alternative haplotype of chimpanzee chr5, homologous to HSA6, approximately 500 kb from the centromere on the p arm (chr5_hap2_hsa6: 62,935,923-63,012,517p; Figure 4A). This is the only NUMT in our dataset longer than the total linear length of chimpanzee mtDNA (16.5 kb). This NUMT comprises multiple mtDNA copies repeated in tandem and non-contiguously (Figure 4B). Multiple (a total of three) tandem copies likely originated due to repeated expansion events, and/or rolling circle amplification of an mtDNA molecule ^48^. The alignable portions of the copies of mtDNA in this NUMT are identical in sequence, indicating this complex NUMT was formed within a relatively short period or has undergone gene conversion across the array. Each mtDNA unit within this NUMT has a >99.23% sequence identity to mtDNA. Assuming a mutation rate of 10^-7^ per site per year in mtDNA ^49^ and 10^-8^ per site per year in the nuclear genome ^50^, we estimate that this NUMT inserted approximately ∼40,000-400,000 years ago.

**Figure 4.**
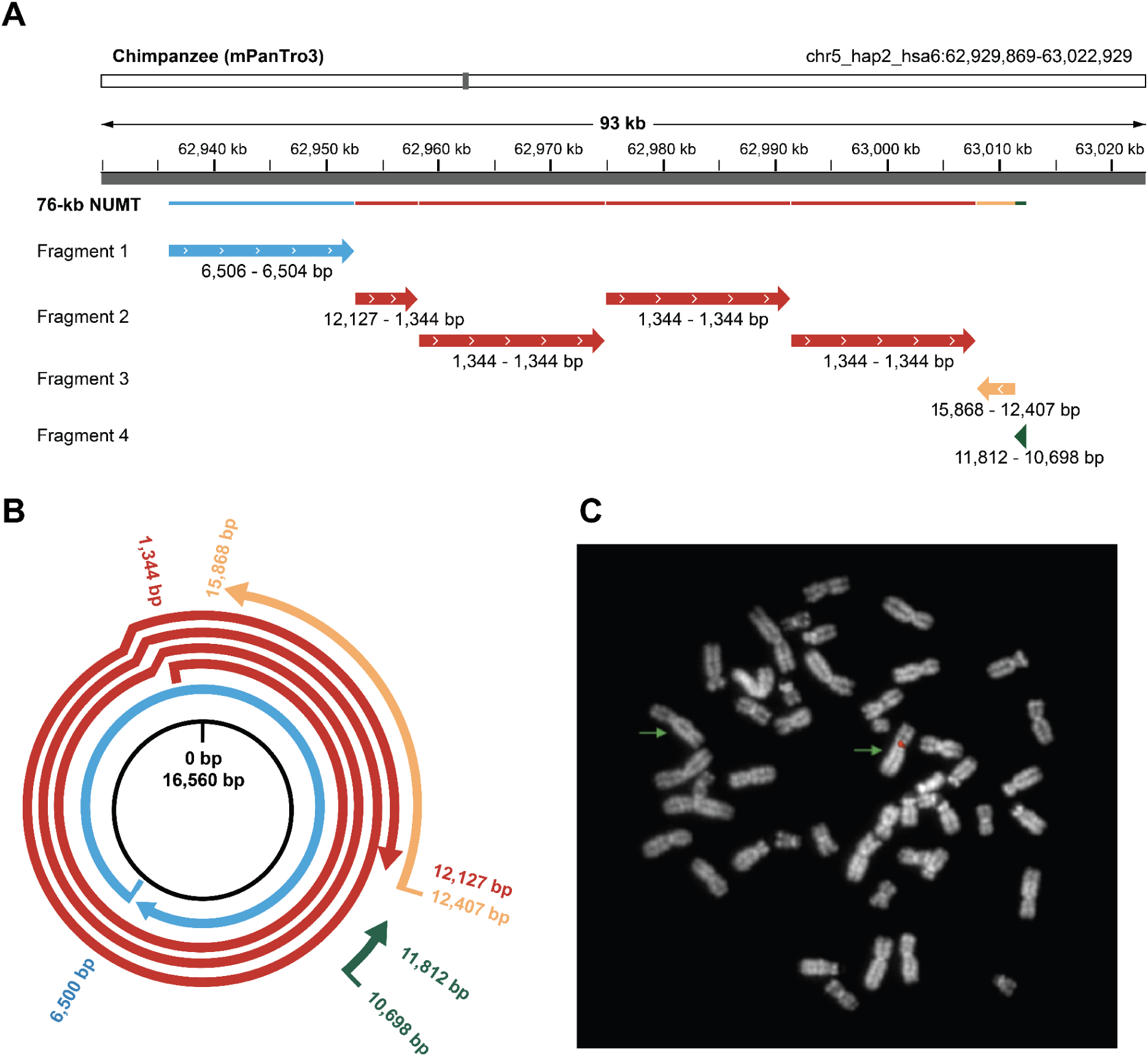
An unusually large, 76-kb NUMT was identified in a chimpanzee assembly. **(A)** Structure of a 76-kb NUMT in the alternate haplotype of chimpanzee chr5 (HSA6), consisting of ∼4.5 copies of mtDNA, visualized in IGV (Integrated Genome Viewer) and modified for clarity. The NUMT was annotated by their homologous coordinates (and strand direction) in mtDNA. Note that full copies homologous to mtDNA start and end at the same coordinate due to the circularity of the genome. These copies were grouped by color to demonstrate contiguity when compared to the circular mtDNA genome, and differ in color to demonstrate when NUMT copies were not contiguous (e.g., copies colored in red span multiple full mtDNA copies and are contiguous to each other in homology to mtDNA). Each copy was >99% identical in sequence to chimpanzee mtDNA, and overlapping sequences within the NUMT array showed no divergence. **(B)** Schematic of NUMT copies and their breakpoints with respect to their homology to chimpanzee mtDNA. The colors correspond to those in A. **(C)** FISH validation of polymorphic NUMT on T2T mPanTro3 chimpanzee chr5 (HSA6). Merged image obtained by overlapping Cy3 fluorochrome signals and DAPI staining. Green arrows indicate the chr5 (HSA6) homologs. Note that both sister chromatids have a pericentromeric hybridization signal in just one of the two homologous chromosomes. PCR amplicons of the NUMT array are included in Figure S8, with the primers detailed in Table S2. FISH results for an unrelated chimpanzee, and a human are included in Figure S9.

We found support for this NUMT array from multiple Oxford Nanopore Technology (ONT) long sequencing reads that span the 76-kb region and from Pacbio HiFi reads that refine, though do not span, the consensus sequence (Figure S7). The region was flagged during assembly validation ^38^ likely due to many reads that align ambiguously to the NUMT region (Figures S9B, S7).

To further validate the presence of this large NUMT, we synthesized fluorescence *in situ* hybridization (FISH) probes using long-fragment PCR (Figure S8) to cover most of the 76-kb region. These probes hybridized with a strong signal to one of the two chr5 (HSA6) in the AG18354 cell line (the original chimpanzee cell line for which the T2T genome was sequenced; Figure 4C) but not to the second homologous chr5 (HSA6) in this same sample, confirming that it is present in only one haplotype. Furthermore, the probes did not hybridize with a strong signal to an unrelated chimpanzee individual nor a human sample (Figure S8).

### NUMT expansion at *Pan* Y-chromosome palindromes

The Y chromosome varies greatly in size and composition between great apes due to its rapid evolution ^41^. In apes, these chromosomes comprise ancestral, pseudoautosomal, satellite, and ampliconic regions (including palindromes), and a human-specific X-transposed region ^41^. We found NUMTs to be absent from ancestral regions and in general rare outside ampliconic regions on chrY, but to be abundant at Y ampliconic regions across all great apes (Figure 5A). However, only the *Pan* genus contained NUMTs within palindromes. This genus displayed a ∼100-fold NUMT enrichment at palindromes compared to non-palindromic ampliconic regions (Figure 5A). These NUMTs were present as paired inverted repeats within palindromes (Figure 5B). They followed a pattern of rapid expansion and gene conversion both within and between palindromes, as evidenced by the high identity of NUMTs, particularly between arms of the same palindrome but also between palindromes (Table S3). The palindrome expansions were evident from self-self global alignments of bonobo and chimpanzee Y chromosomes (Figure 2 in Makova et al. 2024 ^41^).

**Figure 5.**
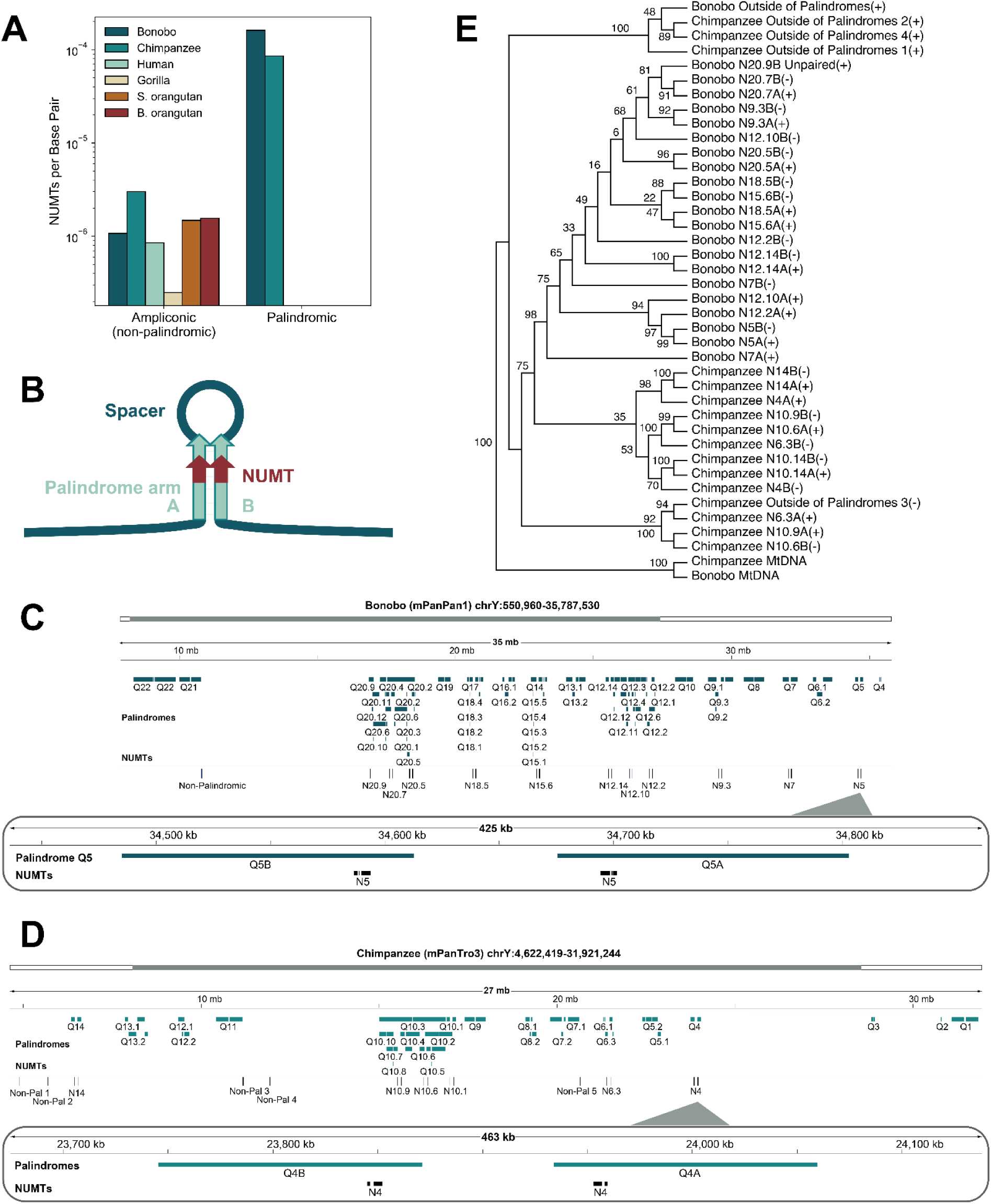
NUMTs in *Pan* Y chromosome palindromes. **(A)** The prevalence of NUMTs per Y chromosome sequence classification. A detailed list of these NUMTs can be found in Table S4. **(B)** Schematic of palindrome structure with NUMTs on each arm. **(C,D)** IGV plot of *Pan* genus lineage-specific expansion of NUMTs in palindromic regions of the Y chromosome, displayed for **(C)** bonobo and **(D)** chimpanzee. These are ∼7-kb NUMTs distributed as pairs along each palindromic arm. Palindromes were annotated with a Q and a number, with each palindrome arm denoted as A and B according to ^41^. Not all palindromes are labeled in these plots, as some overlap. **(E)** An unscaled maximum parsimony tree of palindromic NUMT sequences in bonobo and chimpanzee. We utilized bonobo and chimpanzee mtDNA as outgroups. Branches contain bootstrap values. Branches with <50% support were collapsed. The suffix (+) indicates the NUMT follows the 5’ to 3’ direction, whereas (-) indicates the NUMT follows the 3’ to 5’ direction. PAR: Pseudoautosomal regions.

In bonobo, each NUMT copy in an inverted pair consisted of three regions of sequence identity to mtDNA separated by spacers (homologous to the contiguous positions 13,703-15,438 bp, 15,823-16,054 bp, and 1-4,348 bp in bonobo mtDNA), but overall spanning 7.4 kb (Table S4). Each bonobo NUMT had 69-76% sequence identity to bonobo mtDNA, indicating that these are not recent insertions (Table S3), though the positions with homology to mtDNA suggest a single insertion event within a palindrome arm. For each NUMT pair, one NUMT copy had an inverted NUMT copy on the opposite arm of the same palindrome, separated by ∼100 kb (Figure 5C). These NUMT pairs were observed 11 times in bonobo (located in the palindrome region spanning Y chromosome positions ∼10 to 35 Mb; Figure 5C) along the entire span of palindromic regions. NUMT pairs were highly similar (>99%) between and within palindromes (Table S3), consistent with a rapid expansion. We also observed a non-palindromic NUMT (located at position ∼10 Mb on bonobo chrY) that had a similar sequence to those within palindromes (Figure 5C), suggesting this could be the ancestral sequence to the palindromic NUMTs. This non-palindromic NUMT was separated from palindromic NUMTs by the centromere, and thus could have been part of a deteriorated palindrome.

In chimpanzee, each NUMT copy was 7.1-kb long and included two-three regions of homology to mtDNA separated by a spacer (homologous to the contiguous positions 1-4,289 bp, 13,643-15,375 bp, and 15,666-15,992 bp in bonobo mtDNA; sometimes the second region of homology was missing) on each arm of a palindrome (Table S4). These had between 69-76% sequence identity to chimpanzee mtDNA (Table S3), and were observed 6 times in chimpanzee (located in the palindrome region spanning Y chromosome positions ∼5.5 to 24 Mb) along the entire span of palindromic regions (Figure 5D), similar to bonobo. NUMTs in chimpanzee palindromes were similar in sequence between and within palindromes (Table S3), and we observed four NUMTs outside of palindromes that were similar in sequence to palindromic NUMTs and thus could be the ancestral sequences in chimpanzee.

Considering known homologous palindrome clusters ^41^, these NUMTs were contained in the same ape palindrome cluster, which included multiple palindromes for bonobo (Q5, Q7, Q9, Q12, Q15, Q18, and Q20) and chimpanzee (Q4, Q6, Q10, and Q14). Every bonobo and chimpanzee palindrome in this cluster contained a NUMT on each arm. This palindrome cluster also included human Q7 and *Pongo* Q4 palindromes, but they did not harbor NUMTs.

We built a maximum parsimony tree of palindromic Y-chromosome NUMT sequences in bonobo and chimpanzee, including nearby NUMTs that had a similar sequence but were outside of palindromes and using mtDNA as an outgroup (Figure 5E). We could decipher the phylogeny for this Y-chromosome palindrome cluster using the NUMTs as markers, though it was not fully resolved given the high similarity between palindromes. In most cases, NUMTs present at opposite arms of the same palindrome clustered together, suggesting gene conversion. A bonobo NUMT found in just a single arm of palindrome Q20.9 was nevertheless placed alongside the other palindromic NUMTs. In contrast, for bonobo and chimpanzee, non-palindromic NUMTs were diverged enough to be placed in a separate clade (Figure 5E), likely representing ancestral NUMTs integrated into palindromes.

## Discussion

In this study of ape T2T genomes, we explored NUMT evolution by (1) detailing shared and lineage-specific events, (2) determining the rate of NUMT accumulation among great apes, (3) studying regions of the genome where NUMTs are enriched and depleted, (4) associating NUMT and *Alu* elements, which share a similar expansion mechanism, (5) discovering a rapid NUMT expansion within palindromes of *Pan* chrY, and (6) describing a novel mega-NUMT array in chimpanzee.

### NUMTs among great apes

The majority of NUMTs were expected to be shared among great apes, due to the relatively low NUMT integration rate ^7^. This is consistent with our findings indicating that a high fraction of NUMTs is shared by all great apes and by natural clades, compared to the lower fraction of species-specific NUMTs. The incomplete natural clades, where a single species was missing a NUMT, may be explained by a loss, or mutations, of the NUMTs. Many sparsely populated clades held NUMT phylogenies that could only be explained by multiple loss or integration events (or incomplete lineage sorting), suggesting the rarity of these events or limitations in our ability to detect shared NUMTs. Lineage-specific differences in the number of NUMT events could be due to novel insertions, expansions of pre-existing NUMTs, or degradation of larger NUMTs. In the case of the Y chromosome, the NUMT content increased dramatically in certain species, suggesting that there were either lineage-specific expansions of pre-existing NUMTs, or more recent insertions of large NUMTs, and is consistent with high primate chrY variation (Makova et al. 2024). Though the fraction of the genome covered by NUMTs in orangutans varied greatly at chr7 and chr17, the variation was similar between the species, likely due to incomplete lineage sorting resulting in similar polymorphic sites maintained across species.

The highest rates of NUMT insertions were seen in *Hominini*, particularly in the *Pan* genus, consistent with previous work ^15^. Orangutans had a lower NUMT rate than *Hominini*. The rate of NUMT accumulation in gorilla was lower than in the other species. Gorilla experienced an increase in satellite expansion after divergence from *Hominini* ^38^. As we found that satellites were depleted in NUMTs, this could partially explain slower NUMT accumulation in gorilla. As an alternative biological explanation, gorillas have a polygynous mating pattern, distinguishing them from the rest of great apes. Selection linked to mating patterns could influence the rate of NUMT accumulation. The slow accumulation of NUMTs in gorilla might also be an artifact. The gorilla assembly had the highest gap divergence of all the T2T great ape assemblies ^38^, which could partially explain this slower rate of NUMT accumulation, as the phylogenetic trees are sensitive to the quality of alignments between species.

### Selection against NUMT integrations

Though negative selection is likely acting to limit their deleterious integration into protein-coding sequences and enhancer regions, NUMTs are nevertheless tolerated within introns. The insertion of NUMTs into introns has been observed in other species, such as in honeybees ^18^, but at a much lower frequency (∼3% of NUMTs were found in introns) than observed in great apes (∼30% of NUMTs found in introns), though this could be attributed to lower intron content in honeybees.

NUMT integrations within TEs are likely neutral, given that most TEs are located outside functional regions. TEs were modestly enriched in NUMTs, though we did not detect a significant TE overrepresentation in the longer flanking regions of NUMTs (i.e., on a 50-kb scale analyzed with IWTomics). Our study is likely underpowered due to the small number of NUMTs per species. Given the ubiquity of TEs in ape genomes and their role in genome instability ^47^, the possibility that TEs contribute to NUMT integrations and expansions remains open. This relationship should be further explored as more high-quality non-human ape genomes are becoming available. NFNR regions displayed a high NUMT enrichment. The cause of such enrichment remains an open area of study, with the possibility of a genetic purge, genomic turnover, or insertional preference based on sequence characteristics.

Another study showed that mitochondrial and nuclear DNA stress increase NUMT integration *ex vivo* and *in vitro*, and almost half of these occur within gene regions ^22^. Their results disagree with our observation of (presumably) germline NUMTs located at mostly non-repetitive non-functional regions, suggesting different accumulation patterns without substantial stress. We found a third of NUMTs to be located within gene regions, specifically introns, and not coding regions. NUMTs would likely be more tolerated in somatic cells than the germline, as interrupting coding or regulatory regions would have a smaller impact on the organism, leading to mosaicism with age. Studying multiple tissues within an ape individual would highlight somatic NUMT mosaicism, which has been linked to early mortality ^31^ and cancer ^32^ in humans. In general, we look forward to more high-quality genomes, sequence annotations, and diverse somatic sequencing for non-human primates that would allow us to study evolutionary processes acting on NUMTs in more detail.

### TEs and NUMTs

NUMTs can be seen as remnants of DNA stress. Like TEs, they are expected to integrate in regions of genome instability. We therefore expect a positive correlation between NUMTs and TEs. However, we found a weak negative correlation between NUMT and TE occurrence in the genome, and no significant enrichment in TEs at NUMT 1-50 kb flanks. However, we found an enrichment in TE annotations when considering 50-bp flanks for NUMTs. This may reflect that NUMTs have only local integration preferences close to TEs. A major limitation to understanding NUMT integration preferences is their relatively small number. Our study in particular was limited to a single individual per species.

### A large, heterozygous NUMT in chimpanzee

We discovered a polymorphic NUMT event 76 kb in size—much longer than the other NUMTs—in chimpanzee. The assembly for this region was flagged as potentially erroneous, due to the high number of reads mapping to both mtDNA and chr5 (HSA6). However, we were able to identify ONT Ultra-Long reads that span the entire 76-kb NUMT, and PacBio HiFi reads that align unambiguously to this NUMT (Figure S7). Moreover, we verified the location and heterozygous nature of this NUMT using FISH (the limitations of this approach are described below).

The domestic cat (*Felis catus*) is also known to house a large NUMT near the centromere of chrD2, though older and much larger than the chimpanzee NUMT. The cat NUMT likely inserted ∼2 MYA as a single NUMT, then amplified to range in size from 300-600 kb in the population ^2^. So-called mega-NUMTs have also been observed in human pedigrees, tandemly repeated 56 times the size of a human mtDNA ^51^. If located in the active region of the centromere, these NUMTs have the potential to interfere with centromere activity, leading to cellular senescence ^30^. Though the chimpanzee NUMT array was found within 0.5 Mb of the chr5 (HSA6) centromere, it was not located within satellite sequence.

### NUMT expansion in *Pan* genus Y-chromosome palindromes

We observed a series of NUMTs highly diverged from mtDNA, but almost identical in sequence among themselves, within homologous palindromes of bonobo and chimpanzee but not human or other great apes. Outside of the *Pan* genus, the homologous palindrome was present only once per species (missing in gorilla) but none included the NUMT. Since this NUMT was present in all *Pan* genus palindromes belonging to the homologous cluster, but none outside of *Pan*, we hypothesize that it inserted in the Y chromosome region of the chimpanzee and bonobo common ancestor, within a palindrome arm. Gene conversion likely copied the NUMT to the opposite palindrome arm. Then this NUMT-containing palindrome expanded multiple times leading to the phylogenies observed in bonobo and chimpanzee. This insertion likely occurred within 4.5 MYA of evolution (2.5 to 7 MYA), judging by species divergence estimates. We also expect gene conversion to occur for NUMT sequences between palindromic arms, homogenizing the NUMT sequences between the palindrome arms and also across palindromes. Indeed, the NUMT sequence divergence between palindromic arms is less than 1%. This is consistent with our observation that NUMTs in the same palindrome were often grouped together on the phylogenetic tree and can be explained by gene conversion among palindromes and a rapid expansion of palindromes in the *Pan* genus ^41^. This finding of NUMTs within the Y chromosome shines more light into this enigmatic genomic region.

### Limitations of the present study

We annotated NUMTs for a single individual per species. A recent population study demonstrated a high prevalence of polymorphic NUMTs in humans ^7^. Our work likely misses a large fraction of intraspecific NUMT diversity in non-human primates. Additionally, some NUMTs might have integrated during cell line propagation, as the T2T genomes were sequenced from cell lines ^38^.

Our dataset’s relatively small number of NUMT integrations limited our ability to discern integration preferences for a diverse set of genome annotations. For example, recombination and double-strand break maps are expected to be associated with NUMT integration, but we did not find this signal as a result of our analysis. An enrichment of transposable elements within 5-kb of NUMTs has been demonstrated across mammals ^15^, though we found no significant enrichment in either the immediate NUMT flanks (Figure 3) or within 50 kb of NUMTs (Figure S5). We expect a larger dataset to further our understanding of NUMT integration patterns.

Though we were able to validate the presence of a large and heterozygous NUMT on chimpanzee chr5 (HSA6), we were not able to experimentally validate the number of individual copies it comprises. The assembly suggests that this NUMT has four full mtDNA copies, with some shorter copies in the inverse direction. A droplet digital PCR approach ^52^ of a repeating unit unique to the NUMT may yield a proper validation of the total size. However, the low divergence between the NUMT and the extant chimpanzee mtDNA complicates this. A fiber-FISH approach ^53^ could compensate for this, as the NUMT flanks and the content would be distinguishable from each other.

### Future work

Exploring NUMT sequence evolution could elucidate the history of mutation and expansion post-integration of NUMTs. Now that T2T genome assemblies for great apes are available, and their NUMT content assessed, they open up a new era in population genomics of great apes. The ongoing Human Pangenome Reference Consortium project ^54^ provides high-quality genomes for human populations, allowing for similarly high-quality annotation of NUMTs and their sequence context, as previous studies have depended on *de novo* assembly of NUMTs ^14^. Expanding our analysis to population samples of non-human great apes would benefit our understanding of NUMT evolution in this clade. Further, multiple species of lesser apes would be the logical next step for our analysis. Siamang is the only current gibbon species with a T2T genome, though it aligns unreliably to great ape genomes due to rapid karyotype evolution, requiring more gibbon genomes to resolve the NUMT evolution in lesser apes.

### Significance

This is the first comprehensive study of NUMT insertions in non-human primates with high-quality, T2T genome assemblies. NUMT annotations for T2T genomes will be a valuable tool for mtDNA analysis in great apes. As has been done for humans ^7^, an extensive population-scale study of NUMTs in non-human apes will be important for attaining a more complete picture of their evolution.

## Methods

### Reference genomes

We utilized the recently released version 2 of T2T genomes for bonobo (mPanPan1), chimpanzee (mPanTro3), human (CHM13), gorilla (mGorGor1), Sumatran orangutan (mPonAbe1), and Bornean orangutan (mPonPyg2) ^38,41^. The primary assembly was utilized for our analyses unless noted otherwise. In this paper, we refer to human chromosomes and their homologs (HSA) ^38^, not species-specific chromosome numbering. To compare the human chromosome 2 to homologous chromosomes in the apes, we used the site of fusion, a ∼100 kb region (2q14.1, chr2:113,940,058-114,049,496), that separates the ancestral segments corresponding to HSA2a and HSA2b ^55^.

### Identification of NUMTs

To identify candidate NUMTs, the mitochondrial genome for each species was aligned to the corresponding nuclear genome. We utilized the *blastn* aligner (v2.16.0+) with parameters previously used to study NUMTs in the human CHM13 assembly (-evalue 0.0001 -gapopen 5 -gapextend 2 -penalty −3 -reward 2 -task blastn) ^10^. Alignments longer than 30 bp were considered. Overlapping NUMT annotations were merged using bedtools (v2.31.1) and considered the same NUMT insertion.

### Presence/absence phylogenetic analysis

To identify species-specific, lineage-specific, and shared NUMTs, we followed a methodology similar to that used in Yoo et al. 2025 ^38^ for transposable element integration analysis. In brief, we extracted 500-bp flanking sequences on each side of each NUMT, concatenated these sequences, and aligned them against every ape genome using BLAT. We then compared the 14-bp *k*-mer distributions between the candidate matches and the original NUMT with its flanking sequence. Matches with an identity of ≥50% were retained. NUMTs aligned to a single species were classified as species-specific and counted as a single NUMT integration event, even if observed multiple times within that species (Figure 2A). These data were used to build a maximum parsimony tree using PAUP (v4.0a) ^56^ and plotted using MEGA11 ^57^(Figure 2B). The pairwise distance matrix was estimated in MEGA11 using maximum composite likelihood ^58^, which accounts for multiple hits by computing Tamura-Nei distance ^59^. The NUMT accumulation rate for each branch was estimated by dividing the number of NUMTs shared by that branch by the divergence time (see Figure 1A in Makova et al. 2024 ^41^ and references therein).

### Genome annotations

We utilized functional annotations provided by Mohanty et al. ^44^which were derived from the NCBI Eukaryotic Genome Annotation Pipeline and RepeatMasker annotations published alongside the new ape T2T assemblies ^38,41^. Each genome was divided into protein-coding genes (subdivided into coding sequence (CDS), introns, and 5’ and 3’ UTRs), non-protein-coding genes, and repetitive regions (subdivided into satellites, transposable elements, and low complexity regions). The remaining fraction of the genome was categorized as non-functional non-repetitive regions (NFNR). To compute the number of NUMT events and NUMT base pairs that overlap these genome annotations (Figure 3A), at least 10% of the NUMT was required to overlap the feature. The fraction of functional regions of a certain type covered by NUMTs was calculated by dividing the fraction of NUMTs (or NUMT base pairs) within these regions by the fraction of the genome covered by these regions (Figure 3B).

### Enrichment analysis

To study the distribution of NUMTs along each genome we utilized NUMT flanks (50 bp in length) and at least 80% overlap in one NUMT flank and genome annotations was required (Figures 3, S5). Enrichment was computed as the fraction of total NUMTs observed at a genomic feature, normalized by the fraction of the genome covered by that feature. To obtain 95% confidence intervals, we performed a bootstrap procedure ^60^ by resampling each genome annotation with replacement 1,000 times and re-computing the enrichment score for each sampling.

### Functional Hypothesis Testing

To study potential enrichment in the sequence context of NUMTs, we performed functional hypothesis testing using the IWTomics package ^61^ (Figure S5). The test set consisted of 50-kb flanks around NUMTs concatenated into 100-kb windows. The control set was 100 kb windows in the parts of the genome not included in the test set. We considered the following genome annotations (derived from Mohanty et al. 2024^44^) that may be related to NUMT accumulation: origins of replication; CpG islands, non-B DNA motifs ^62^: A-phased repeats (APR), G-quadruplexes (GQ), mirror repeats (MR), and Z-DNA (Z); transposable elements ^38^: short interspersed nuclear elements (SINEs), long interspersed nuclear elements (LINE1, and LINE2/3), long tandem repeat retrotransposons (LTRs), DNA transposons, retroposons, and rolling circle transposons; and other repeat annotations ^38^: satellites, simple repeats and low complexity regions.

### Genome-wide comparison of NUMTs and TEs

To compare NUMT and TE densities across the genome, we computed such densities among non-overlapping genome windows (0.1 Mb and 1 Mb in length) for each species. The genome windows were created using *bedtools makewindows* and the density was computed using *bedtools annotate*. A Pearson correlation test was utilized to compare NUMT density (in log scale) to TE density.

### Long-fragment Polymerase Chain Reaction (PCR)

Long-fragment PCR to make probes for validating the long NUMT on the *Pan* chrY was performed following the manufacturer’s instructions (Vazyme, 2x Phanta Max Master Mix Version 23.1) with minor modifications. Briefly, PCR reactions were performed with 2x Phanta Max Master Mix 25 μl, primers forward and reverse (10 μM) 2 μl each, and 50 ng of DNA adding ddH2O up to 50 μl. The reaction was performed under these conditions: (1) initial denaturation 95°C for 3min, 35 cycles of steps (2) denaturation 95°C for 15 seconds, (3) annealing at 60°C for 15 seconds, and (4) extension 72°C for 16min and a final elongation step at 72°C for 5min.

### Fluorescence *in situ* hybridization (FISH)

The metaphase spreads were obtained from lymphoblast cell lines with standard procedures ^63^) from a chimpanzee cell line (GM18354), a chimpanzee (PTR17, *Pan troglodytes*), and a human HapMap individual (GM18548, Coriell Cell Repository, Camden, NJ, USA) available at the University of Bari. Cytogenetic analyses were performed by fixing metaphase spreads onto slides and then incubating at 90°C for 1 h 30 min to contribute to sample fixation and dehydration. The 0.005% pepsin/HCl 0.01 M treatments were performed to eliminate cytoplasm proteins for better hybridization rates. Subsequent treatments in 1×PBS, MgCl_2_ 0.5 M, 8% paraformaldehyde, and 70%/90%/absolute alcohols allowed proper stabilization, fixation, and dehydration of metaphases and nuclei DNA molecules. FISH experiments were performed as previously described ^64^: 200 ng of the DNA probe (originating from long PCR fragments above), labeled by nick-translation with Cy3-dUTP or fluorescein-dUTP, were precipitated by ion-exchange alcohol precipitation with canine Cot1 DNA and finally denatured for 2 min at 70°C and hybridized at 37°C overnight. Post-hybridization washing was done at 60°C in 0.1× SSC (three times, high stringency). The slides were then stained with DAPI, producing a Q-banding pattern. The fluorescence signals from Cy3, Fluorescein, and DAPI were detected separately with specific filters using a Leica DMRXA epifluorescence microscope equipped with a cooled CCD camera (Teledyne Princeton Instruments, Acton, MA, USA) and recorded as grayscale images. A total of 50 cells per experiment were scored. Finally, the Adobe Photoshop™ software (2024) was used for image pseudo-colorization and merging of the acquired images.

### Time of divergence

We utilized the following formula to estimate the time of divergence for NUMTs: *T = d/2u*, where *d* is the divergence between sequences and *u* is the mutation rate. Given the uncertain time of insertion of the NUMT, we were conservative in estimating time of divergence by assuming mutation rates of 10^-7^ substitutions per site per year for mtDNA ^49^ and 10^-8^ substitutions per site per year for the nuclear genome ^50^.

## Resource availability

NUMT annotations and how they are shared across the great apes were provided as Supplemental Data Files 1 and 2. The in-house scripts generated for this paper are available on GitHub: https://github.com/makovalab-psu/T2T_greatApe_NUMTs.

## Acknowledgments

We are grateful to Mark Loftus, Miriam K. Konkel, Erin Molloy, Christine R. Beck, Arang Rhie, Gabrielle Hartley, Glennis Logsdon, Prajna Hebbar, and the other members of the T2T Transposable Element and CenSat working groups, for their invaluable feedback in the development of this project. We also thank Saswat Mohanty, Karol Pál, Linnéa Smeds, and Alexis Santos for their critical feedback and discussions that helped improve this project. This work was funded, in part, by the NIH award R35GM151945 and the Willaman Endowed Chair Fund for K.D.M., and the GlaxoSmithKline (GSK) Graduate Fellowship for E.T.G.

## Author contributions

E.T.G.: Conceptualization, Data Curation, Formal analysis, Funding acquisition, Investigation, Methodology, Project administration, Software, Resources, Visualization, Writing - original draft. M.A.C.: Methodology, Software (IWTomics implementation), Writing - review & editing. J.M.S.: Methodology, Resources (Repeat annotations), Writing - review & editing. M.V.: Investigation, Validation (FISH experiment), Writing - review & editing. R.J.O.: Supervision, Funding acquisition, Writing - review & editing. K.D.M.: Conceptualization, Supervision, Funding acquisition, Project administration, Resources, Writing - original draft.

## Supplemental Files

Supplemental files include Supplemental Figures, Supplemental Tables, and Supplemental Data Files 1 and 2.

