## Supplemental Figures for "Nuclear Mitochondrial Sequences in Great Ape Telomere-to-Telomere Genomes"

### Supplementary Figures

**Figure S1. NUMT length (bp) and % sequence identity to mtDNA.**

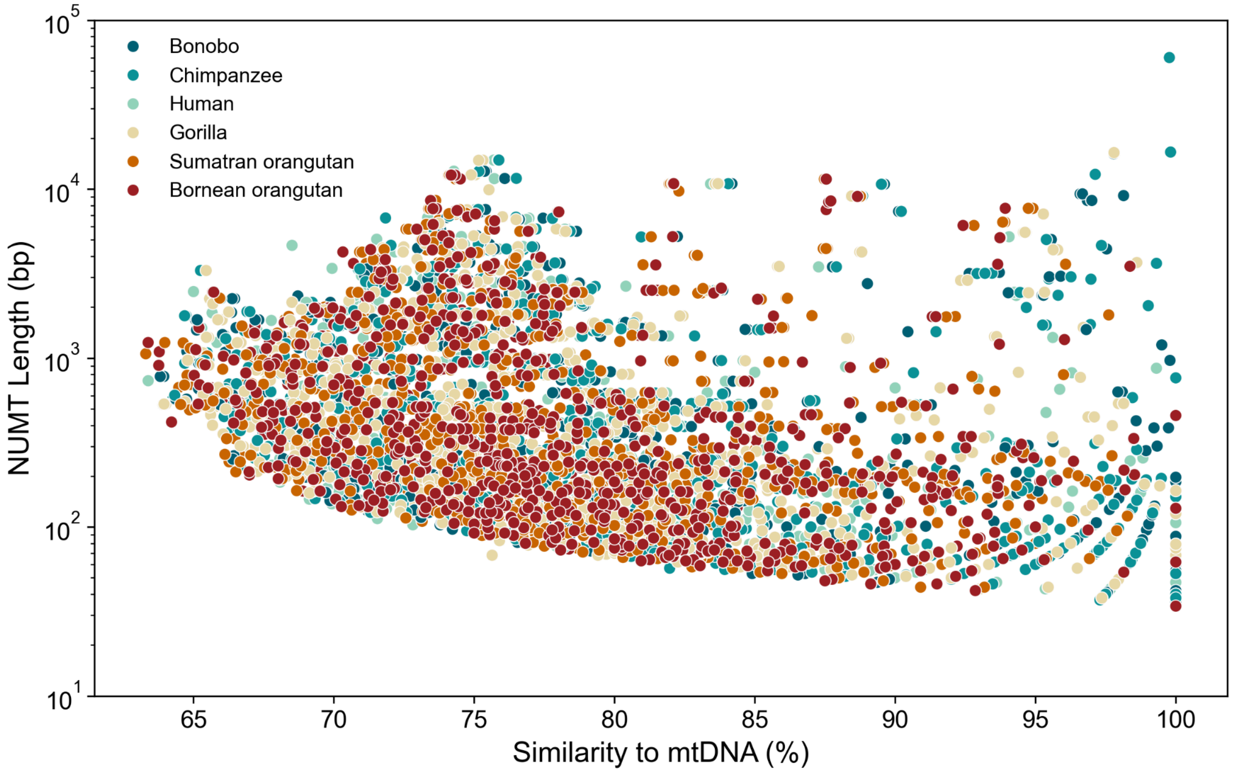

**Figure S2. NUMT content across homologous ape chromosomes (HSA) in alternate haplotype assemblies.** (**A**) Proportion of NUMT lengths per species. (**B**) The number of NUMTs per chromosome per species. (**C**) Total NUMT kilobases per chromosome per species (NUMT content). (**D**) NUMTs per bp for each chromosome and species. (**E**) The fraction of total NUMT length for each chromosome and species. The sex chromosomes included here are the same as in Figure 1 but human CHM13 is a haploid assembly and therefore was not included in this figure.

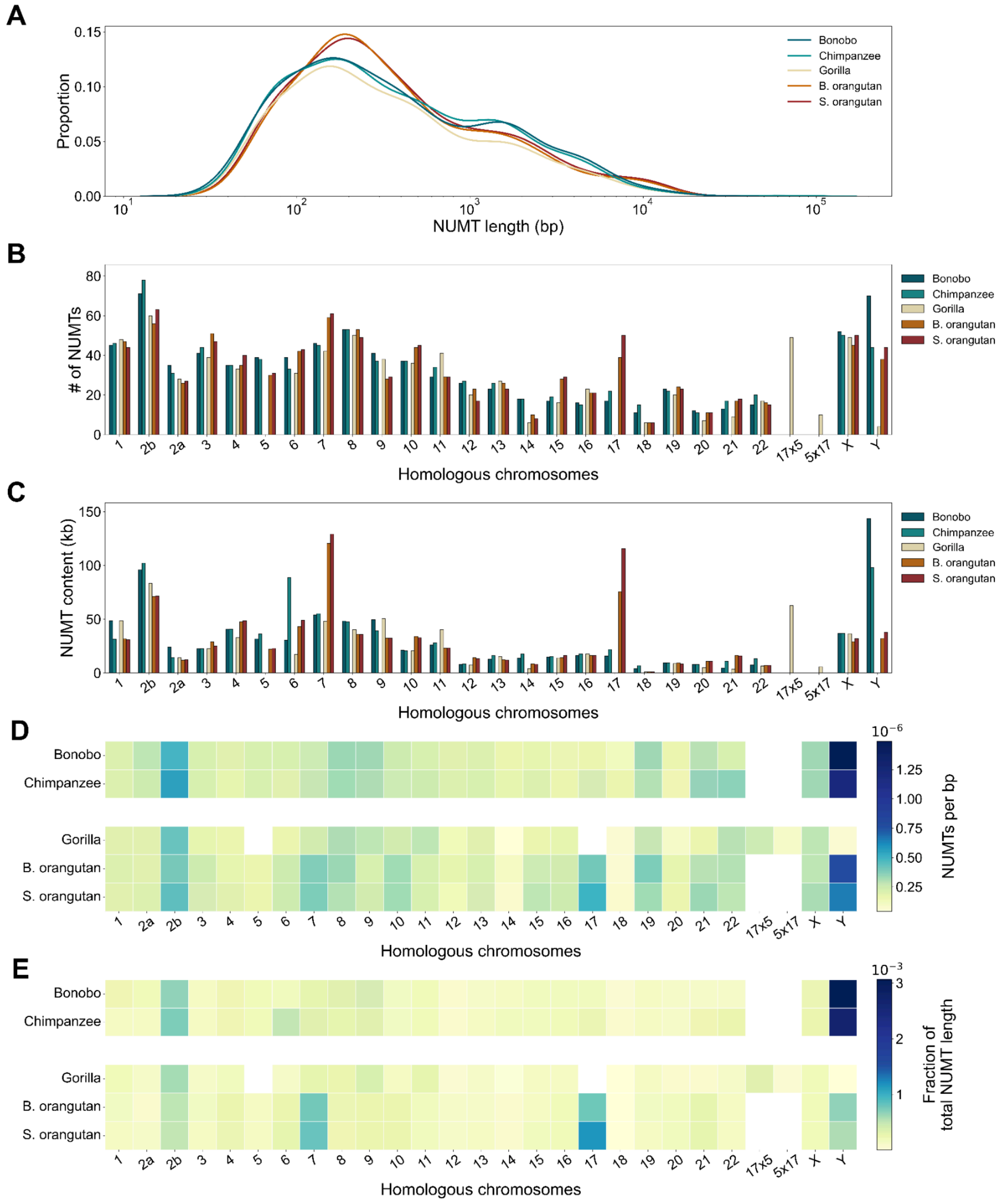

**Figure S3. Shared and lineage-specific NUMTs.** Identical to Figure 2A but includes all clades instead of just the 30 most numerous ones.

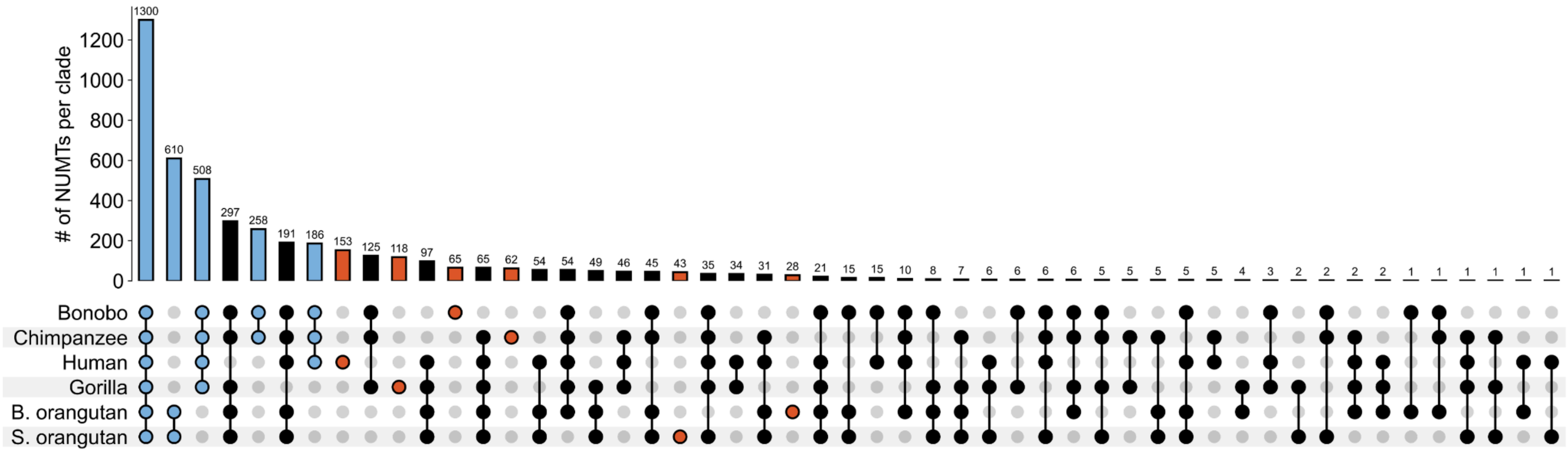

**Figure S4. NUMT enrichment in transposable elements (TEs) varying by the flank size considered.** (A) The percentage of NUMTs flanked by a transposable element. (B) NUMT enrichment in TEs depending on the flank size.

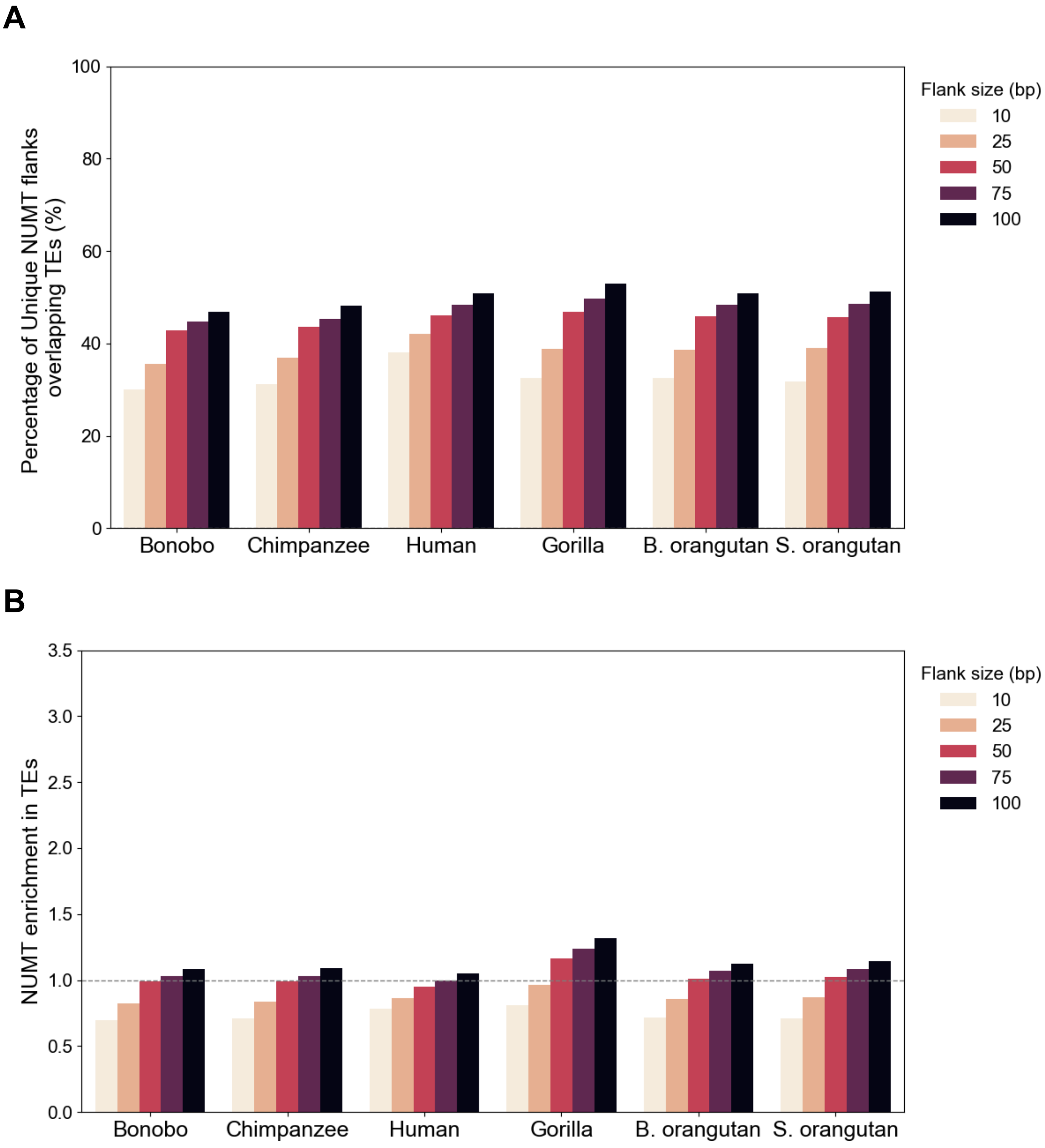

**Figure S5. IWTomics results for NUMT flanking regions versus control regions in human.** These represent 50-kb flanks, split into 1-kb increments, up- and downstream of NUMTs compared to similarly sized and non-overlapping genome windows that exclude NUMTs, used as controls. The Y axis summarizes the results for various genome annotations (see Methods). We used means as the test statistic. The scale of windows on the X axis is in megabases (Mb) upstream and downstream from the NUMT integration point in the middle.

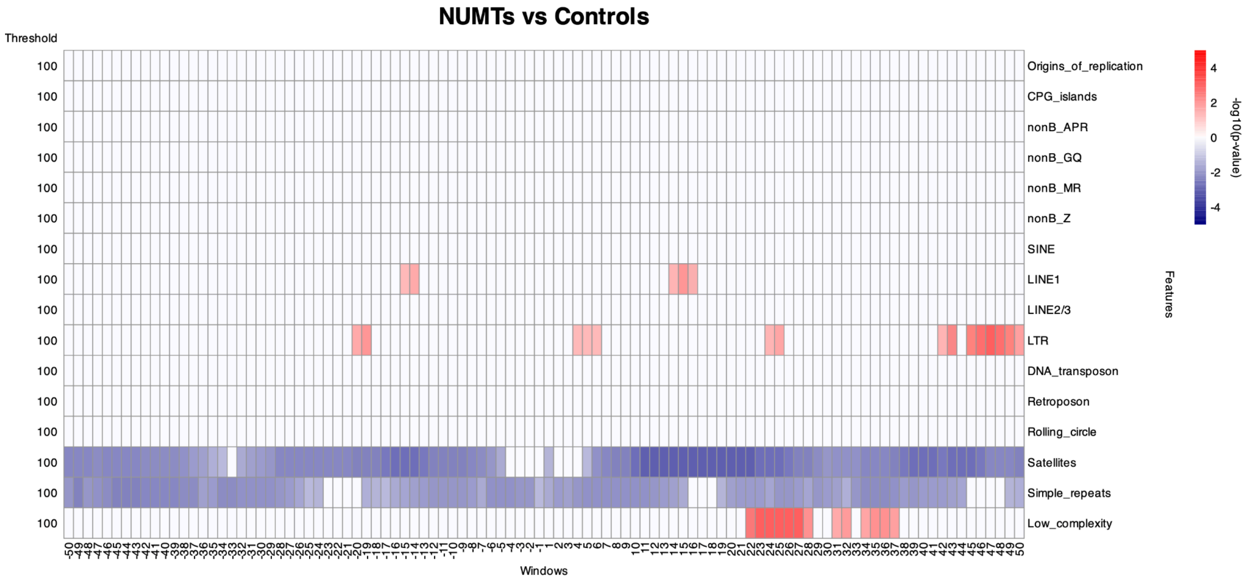

**Figure S6. NUMT and TE density in non-overlapping genomic windows for each species, using 0.1-Mb and 1-Mb window sizes.** These variables were tested using a Pearson's correlation test.

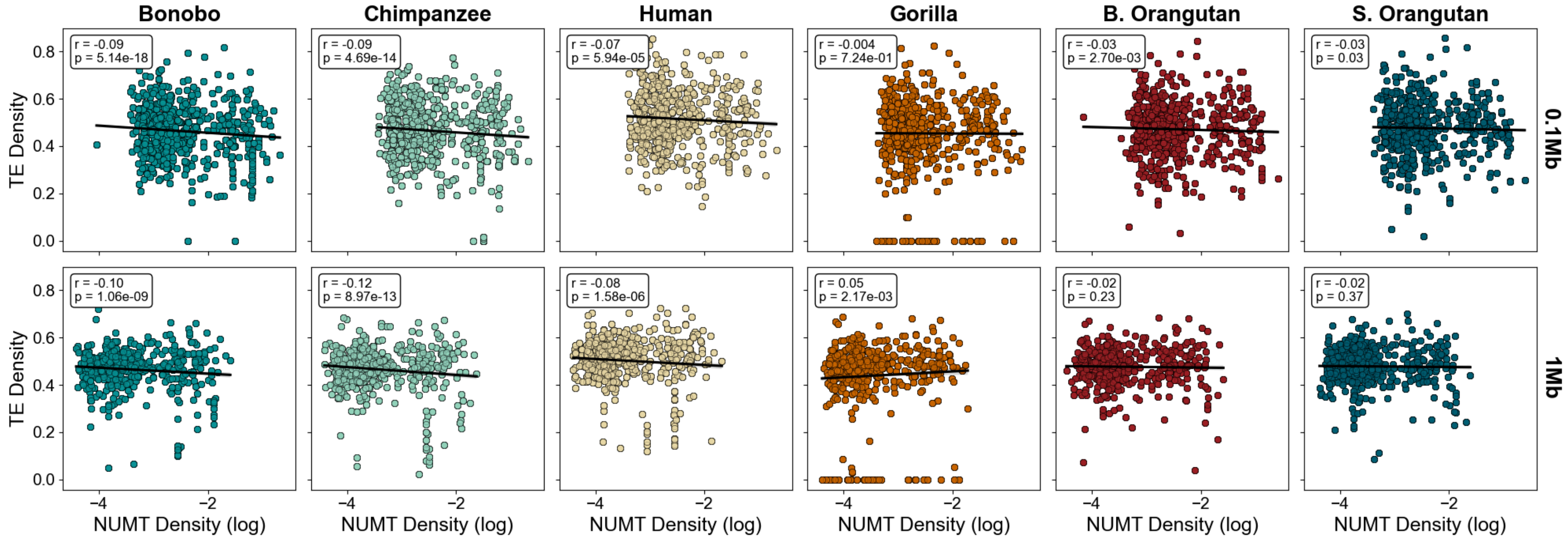

**Figure S7. Long-read DNA sequencing reads utilized to build the mPanTro3 chr5 alternate assembly at the site of a 76-kb NUMT. (A**) IGV track of the 76-kb NUMT array, copies within the array were colored to reflect contiguous homology to chimpanzee mtDNA. This figure is identical to Figure 4 and was included here for convenience. (**B**) PacBio HiFi and (**C**) Oxford Nanopore Ultra-Long reads that were aligned to the NUMT assembly. Aligned reads in *gray* represent and reads in *white* represent secondary alignments. Note the higher read depth in the NUMT compared to the flanking regions (~10 reads/bp). (**D**) Sequencing reads that span the borders between NUMT copies 1 and 2, (**E**) as well as the border between NUMT copies 2 and 3. These alignments only include reads that were primarily aligned to the NUMT. Note that many reads do not span the borders between copies, align to mtDNA, and are ~16.5-kb in length; these represent lower quality alignments. Also note reads that do span the borders between copies (varying in length 38-kb to 440-kb) and do not align simultaneously to mtDNA, supporting the NUMT assembly.

**A**

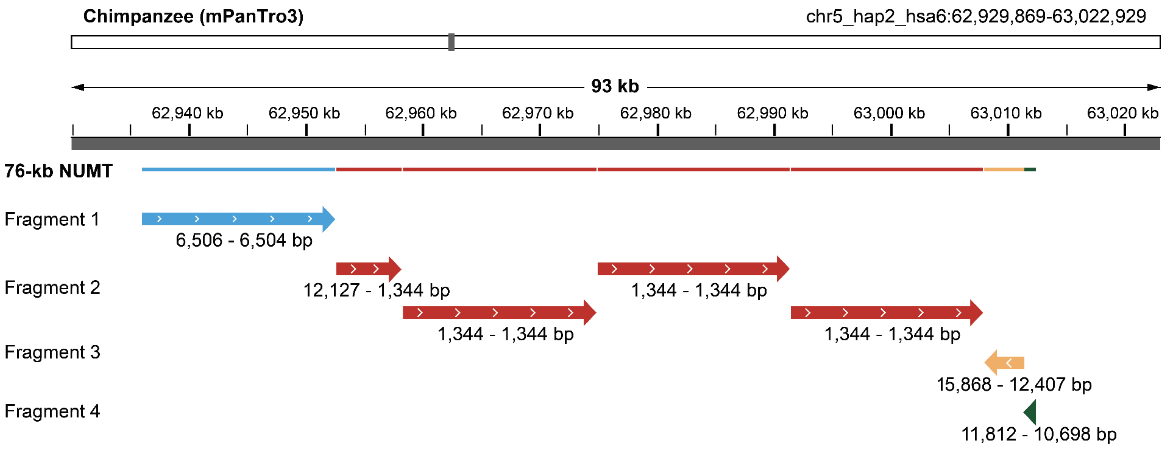

B

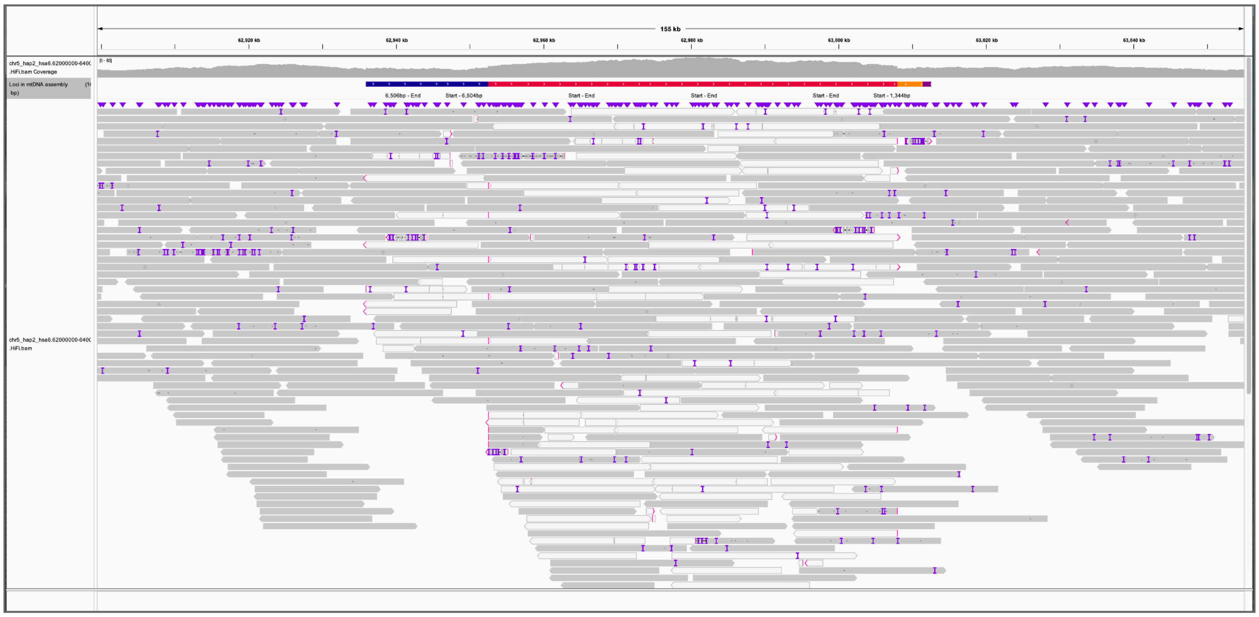
155 kb
62,920 kb
62,940 kb
62,960 kb
62,980 kb
63,000 kb
63,020 kb
63,040 kb
chr5_hap2_hsa6.62000000-640( .HiFi.bam Coverage
Loci in mtDNA assembly bp)
6,506bp - End
Start - 6,504bp
Start - End
Start - End
Start - End
Start - 1,344bp
chr5_hap2_hsa6.62000000-640( .HiFi.bam

C

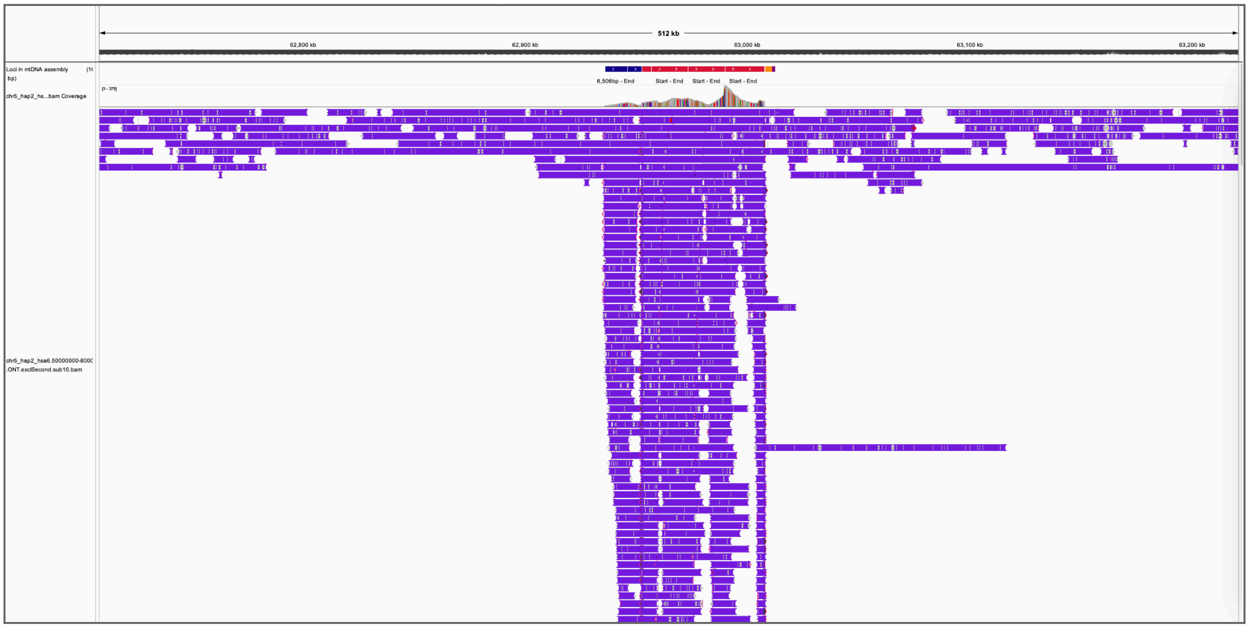
512 kb
62,800 kb
62,900 kb
63,000 kb
63,100 kb
63,200 kb
Loci in mtDNA assembly bp)
chr5_hap2_hs...bam Coverage
6,506bp - End
Start - End
Start - End
Start - End
chr5_hap2_hsa6.50000000-800( .ONT.exclSecond.sub10.bam

D

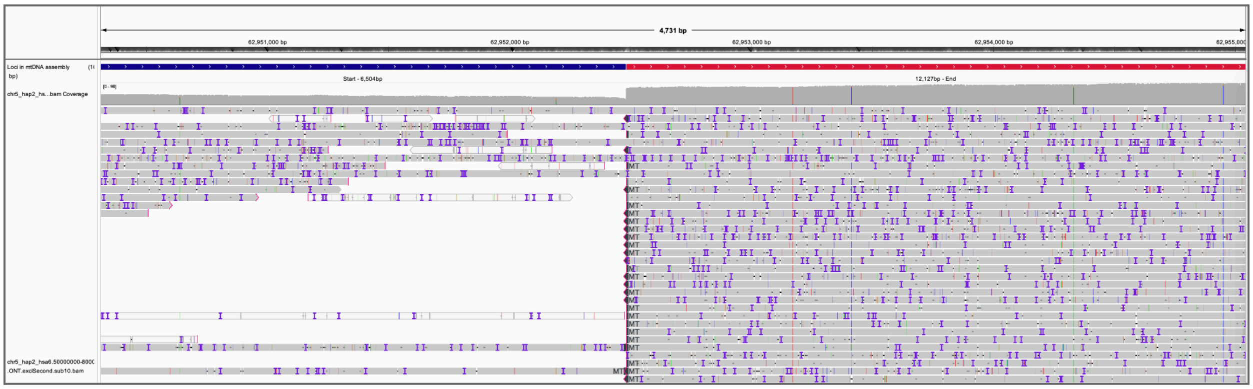
4,731 bp
62,951,000 bp
62,952,000 bp
62,953,000 bp
62,954,000 bp
Loci in mtDNA assembly
chr5_hap2_hs...bam Coverage
Start - 6,504bp
12,127bp - End
chr5_hap2_hsa6.50000000-6000
.ONT.exclSecond.sub10.bam

E

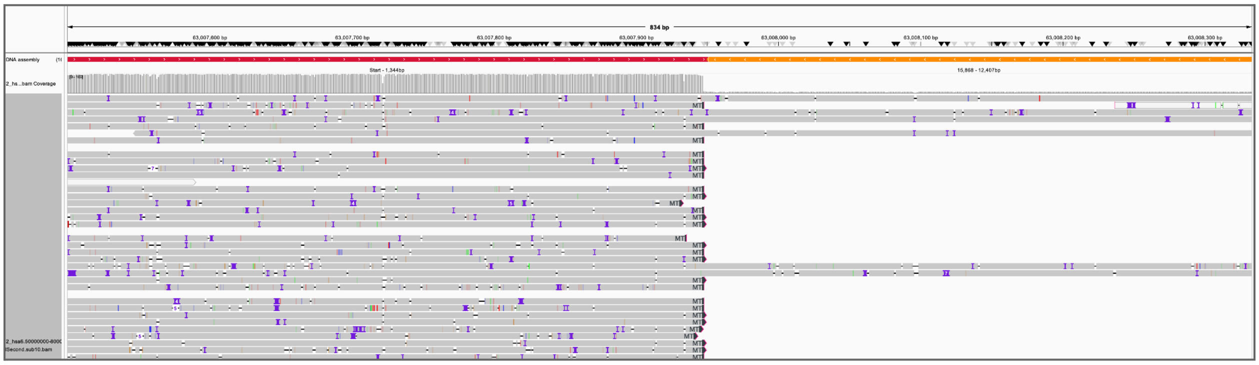
834 bp
63,007,600 bp
63,007,700 bp
63,007,800 bp
63,007,900 bp
63,008,000 bp
63,008,100 bp
63,008,200 bp
63,008,300 bp
Start - 1,344bp
15,868 - 12,407bp

**Figure S8. PCR amplicons targeting the 76-kb NUMT array in chimpanzee. (A)** Gel electrophoresis of long PCR amplicons of NUMT A, B, C, and D on chimpanzee chromosome 5. The lanes contain: (1) GM18354 (T2T chimpanzee); (2) PTR17 (non-T2T chimpanzee); (3) GM18548 (human) and (4) no DNA. The Lambda III DNA ladder was used for scale. The primer design is detailed in Table S3. **(B)** Primers aligned to chimpanzee mtDNA. Clockwise from top left: primer set A, B, C, and D.

A

B

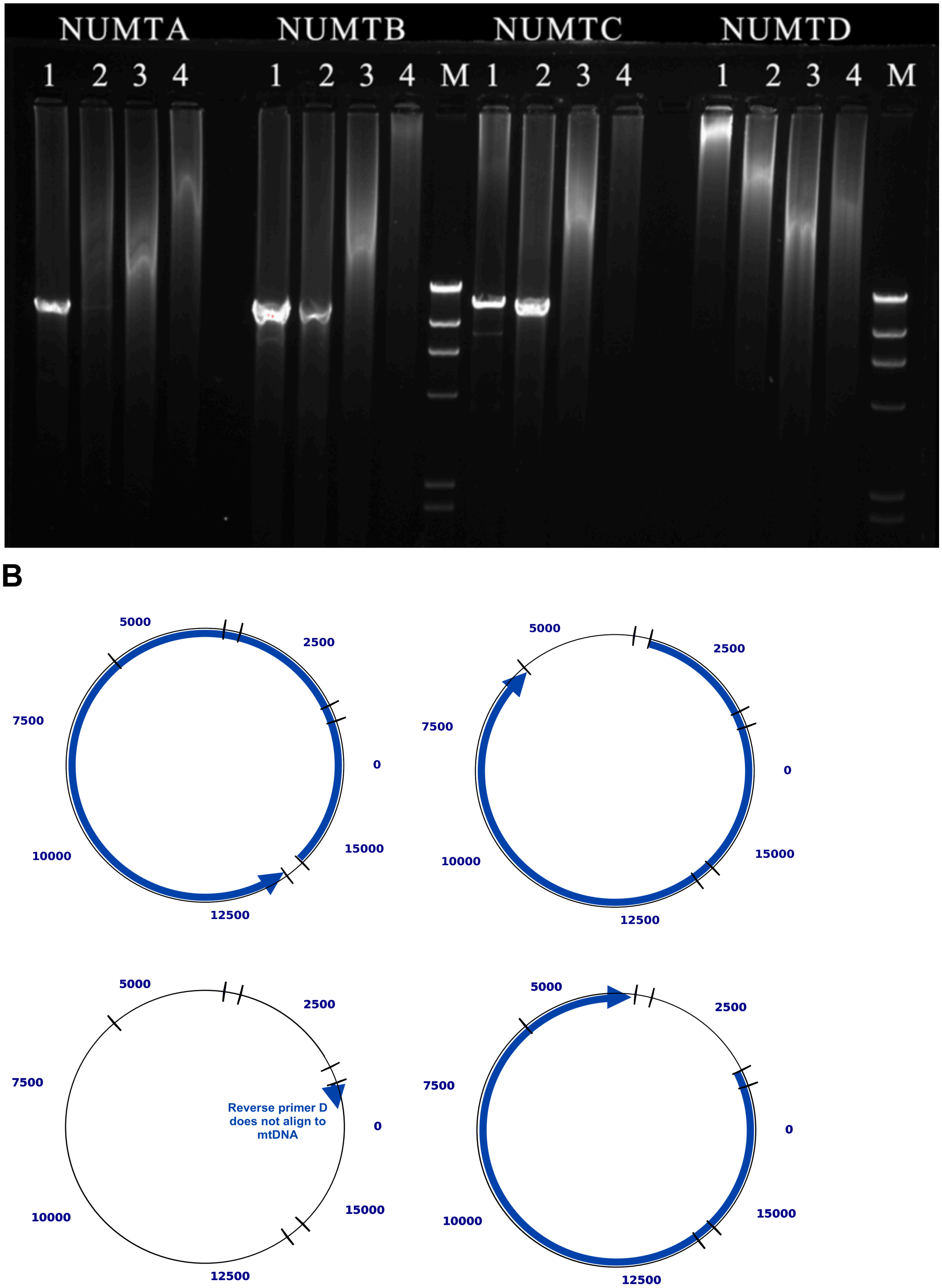
NUMTA
NUMTB
NUMTC
NUMTD
1 2 3 4 1 2 3 4 M 1 2 3 4 1 2 3 4 M
5000
2500
7500
0
10000
15000
12500
Reverse primer D does not align to mtDNA

**Figure S9. FISH probes on other individuals, with no signal of hybridization to the 76-kb NUMT observed in the T2T chimpanzee. (A)** T2T chimpanzee (AG18354): (left) image captured using Cy3 fluorochrome and (right) merged image obtained overlapping Cy3 signals and DAPI staining. **(B)** PTR17 chimpanzee: (left) image captured using Cy3 fluorochrome and (right) merged image obtained overlapping Cy3 signals and DAPI staining. **(C)** GM18548 human: (left) image captured using Cy3 fluorochrome and (right) merged image obtained overlapping Cy3 signals and DAPI staining.

**A**

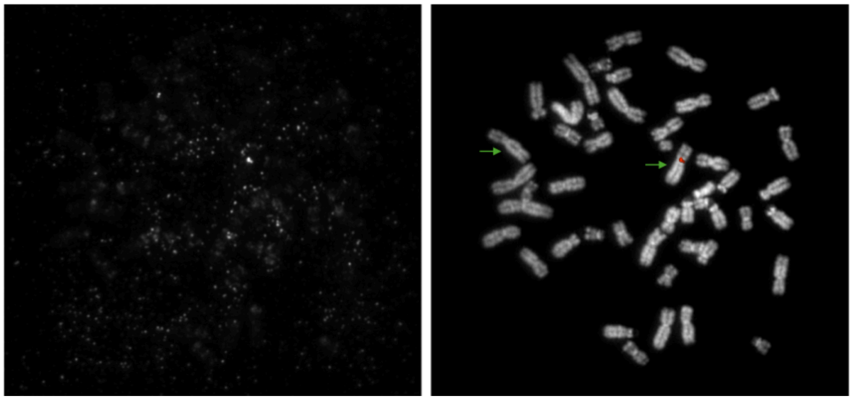

**B**

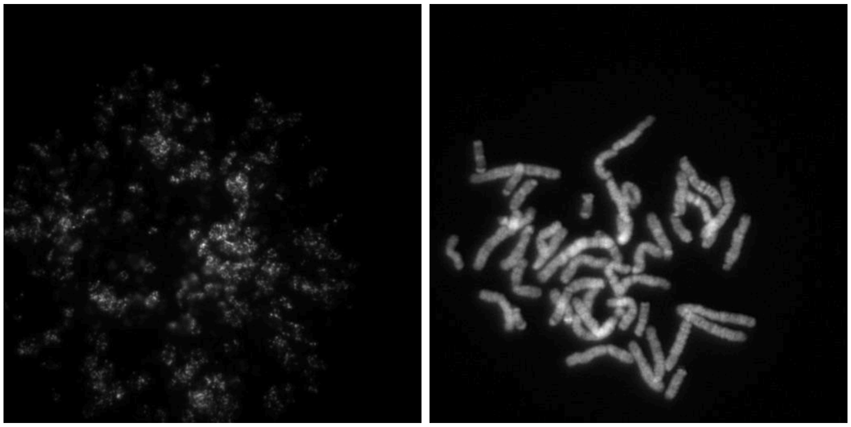

**C**

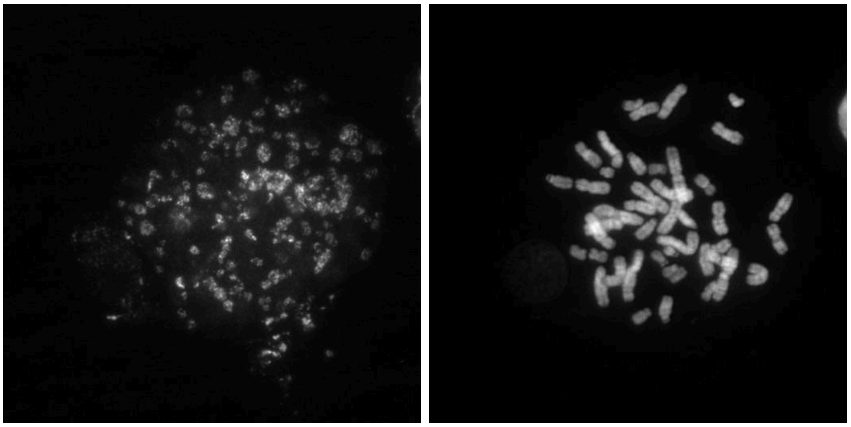
