## Supplemental Tables for "Nuclear Mitochondrial Sequences in Great Ape Telomere-to-Telomere Genomes"

**Table S1. Number of NUMTs per species and haplotype.** The human CHM13 reference is haploid, therefore we only include one haplotype.

| Species | Primary | Alternate |
|---|---|---|
| Bonobo | 713 | 702 |
| Chimpanzee | 717 | 723 |
| Human | 711 | - |
| Gorilla | 648 | 656 |
| Bornean orangutan | 714 | 721 |
| Sumatran orangutan | 715 | 729 |

**Table S2. Primers designed to validate the large NUMT on chimpanzee (mPanTro3) chr5_hap2_hsa6:62935924-63012517**. An IGV image of the primer alignments to the NUMT and to mtDNA are included in Figure 4. A gel electrophoresis image of the amplification products is included in Figure S7.

| Target NUMT start | Target NUMT end | Primer names | Sequences | Melting temperature (ºC) | Product size | Aligns to mtDNA with 100% identity |
|---|---|---|---|---|---|---|
| 1 | 19125 | NUMTA_F | ACAGCACTTCCTTGGCCTAT | 68 | 16196 | 353-376 (+) |
| | | NUMTA_R | AATAGGCTCGGGTGTCTACG | 70 | | 6219-6239 (-) |
| 19126 | 38025 | NUMTB_F | GCTCAGCCTATATACCGCCA | 70 | 14032 | 670-690 (+) |
| | | NUMTB_R | ATCGTGTAAGGGTAGGGCTG | 70 | | 14681-14701 (-) |
| 38026 | 57076 | NUMTC_F | CTCTACATCACCGCCCCAA | 70 | 14000 | 2937-2956 (+) |
| | | NUMTC_R | ATCTAAAACACTCTTTACGCCGG | 70 | | 353-376 (-) |
| 57077 | 76596 | NUMTD_F | AACATGACCCCTGGCCATAA | 68 | 14650 | 3260-3280 (+) |
| | | NUMTD_R | TGCTACTCTCTGGTTTCGGG | 70 | | None |

**Table S3. Tamura-Nei Pairwise Distance matrix of bonobo and chimpanzee NUMTs in chrY palindromic regions.** These are compared to the corresponding mitochondrial genomes. Intensity of the color corresponds to the magnitude of pairwise distances.

| | | 1 | 2 | 3 | 4 | 5 | 6 | 7 | 8 | 9 | 10 | 11 | 12 | 13 | 14 | 15 | 16 | 17 | 18 | 19 | 20 | 21 | 22 | 23 | 24 | 25 | 26 | 27 | 28 | 29 | 30 | 31 | 32 | 33 | 34 | 35 | 36 | 37 | 38 | 39 | 40 |
|---|---|---|---|---|---|---|---|---|---|---|---|---|---|---|---|---|---|---|---|---|---|---|---|---|---|---|---|---|---|---|---|---|---|---|---|---|---|---|---|---|---|
| 1 | Bonobo_Not_in_palindrome_I | 0.000 | | | | | | | | | | | | | | | | | | | | | | | | | | | | | | | | | | | | | | | |
| 2 | Bonobo_Q20.9B_alone_3-nu | 0.133 | 0.000 | | | | | | | | | | | | | | | | | | | | | | | | | | | | | | | | | | | | | | |
| 3 | Bonobo_Q20.7B(-) | 0.133 | 0.001 | 0.000 | | | | | | | | | | | | | | | | | | | | | | | | | | | | | | | | | | | | | |
| 4 | Bonobo_Q20.7A(+) | 0.133 | 0.001 | 0.000 | 0.000 | | | | | | | | | | | | | | | | | | | | | | | | | | | | | | | | | | | | |
| 5 | Bonobo_Q20.5B(-) | 0.132 | 0.002 | 0.001 | 0.001 | 0.000 | | | | | | | | | | | | | | | | | | | | | | | | | | | | | | | | | | | |
| 6 | Bonobo_Q20.5A(+) | 0.132 | 0.002 | 0.001 | 0.001 | 0.000 | 0.000 | | | | | | | | | | | | | | | | | | | | | | | | | | | | | | | | | | |
| 7 | Bonobo_Q18.5B(-) | 0.132 | 0.001 | 0.001 | 0.001 | 0.001 | 0.001 | 0.000 | | | | | | | | | | | | | | | | | | | | | | | | | | | | | | | | | |
| 8 | Bonobo_Q18.5A(+) | 0.133 | 0.001 | 0.001 | 0.001 | 0.001 | 0.001 | 0.001 | 0.000 | | | | | | | | | | | | | | | | | | | | | | | | | | | | | | | | |
| 9 | Bonobo_Q15.6B(-) | 0.132 | 0.001 | 0.001 | 0.001 | 0.001 | 0.001 | 0.000 | 0.001 | 0.000 | | | | | | | | | | | | | | | | | | | | | | | | | | | | | | | |
| 10 | Bonobo_Q15.6A(+) | 0.133 | 0.002 | 0.001 | 0.001 | 0.001 | 0.001 | 0.000 | 0.000 | 0.000 | 0.000 | | | | | | | | | | | | | | | | | | | | | | | | | | | | | | |
| 11 | Bonobo_Q12.14B(-) | 0.133 | 0.003 | 0.003 | 0.003 | 0.002 | 0.002 | 0.002 | 0.002 | 0.002 | 0.002 | 0.000 | | | | | | | | | | | | | | | | | | | | | | | | | | | | | |
| 12 | Bonobo_Q12.14A(+) | 0.133 | 0.003 | 0.003 | 0.003 | 0.002 | 0.002 | 0.002 | 0.002 | 0.002 | 0.002 | 0.000 | 0.000 | | | | | | | | | | | | | | | | | | | | | | | | | | | | |
| 13 | Bonobo_Q12.10B(-) | 0.133 | 0.001 | 0.001 | 0.001 | 0.001 | 0.001 | 0.001 | 0.001 | 0.001 | 0.001 | 0.002 | 0.002 | 0.000 | | | | | | | | | | | | | | | | | | | | | | | | | | | |
| 14 | Bonobo_Q12.10A(+) | 0.134 | 0.004 | 0.003 | 0.004 | 0.003 | 0.003 | 0.003 | 0.003 | 0.003 | 0.003 | 0.003 | 0.003 | 0.003 | 0.000 | | | | | | | | | | | | | | | | | | | | | | | | | | |
| 15 | Bonobo_Q12.2B(-) | 0.132 | 0.001 | 0.001 | 0.001 | 0.001 | 0.001 | 0.000 | 0.000 | 0.000 | 0.001 | 0.002 | 0.002 | 0.001 | 0.003 | 0.000 | | | | | | | | | | | | | | | | | | | | | | | | | |
| 16 | Bonobo_Q12.2A(+) | 0.134 | 0.004 | 0.003 | 0.004 | 0.003 | 0.003 | 0.003 | 0.003 | 0.003 | 0.003 | 0.003 | 0.003 | 0.003 | 0.002 | 0.003 | 0.000 | | | | | | | | | | | | | | | | | | | | | | | | |
| 17 | Bonobo_Q9.3B(-) | 0.133 | 0.001 | 0.001 | 0.001 | 0.001 | 0.001 | 0.001 | 0.001 | 0.001 | 0.001 | 0.003 | 0.003 | 0.001 | 0.004 | 0.001 | 0.003 | 0.000 | | | | | | | | | | | | | | | | | | | | | | | |
| 18 | Bonobo_Q9.3A(+) | 0.133 | 0.001 | 0.001 | 0.001 | 0.001 | 0.001 | 0.001 | 0.000 | 0.001 | 0.001 | 0.002 | 0.002 | 0.001 | 0.004 | 0.001 | 0.003 | 0.000 | 0.000 | | | | | | | | | | | | | | | | | | | | | | |
| 19 | Bonobo_Q7B(-) | 0.132 | 0.002 | 0.002 | 0.002 | 0.001 | 0.001 | 0.001 | 0.001 | 0.001 | 0.001 | 0.002 | 0.002 | 0.001 | 0.003 | 0.001 | 0.003 | 0.002 | 0.002 | 0.000 | | | | | | | | | | | | | | | | | | | | | |
| 20 | Bonobo_Q7A(+) | 0.133 | 0.004 | 0.003 | 0.004 | 0.003 | 0.003 | 0.003 | 0.003 | 0.003 | 0.003 | 0.004 | 0.004 | 0.003 | 0.004 | 0.003 | 0.004 | 0.004 | 0.004 | 0.003 | 0.000 | | | | | | | | | | | | | | | | | | | | |
| 21 | Bonobo_Q5B(-) | 0.134 | 0.004 | 0.004 | 0.004 | 0.003 | 0.003 | 0.003 | 0.003 | 0.003 | 0.003 | 0.003 | 0.003 | 0.003 | 0.002 | 0.003 | 0.001 | 0.004 | 0.004 | 0.003 | 0.004 | 0.000 | | | | | | | | | | | | | | | | | | | |
| 22 | Bonobo_Q5A(+) | 0.134 | 0.004 | 0.004 | 0.004 | 0.003 | 0.003 | 0.003 | 0.003 | 0.003 | 0.003 | 0.003 | 0.003 | 0.003 | 0.002 | 0.003 | 0.001 | 0.004 | 0.004 | 0.003 | 0.004 | 0.000 | 0.000 | | | | | | | | | | | | | | | | | | |
| 23 | Chimpanzee_1_Not_in_pals_ | 0.004 | 0.133 | 0.133 | 0.133 | 0.133 | 0.133 | 0.133 | 0.133 | 0.133 | 0.133 | 0.133 | 0.133 | 0.133 | 0.134 | 0.133 | 0.134 | 0.133 | 0.133 | 0.133 | 0.134 | 0.134 | 0.134 | 0.000 | | | | | | | | | | | | | | | | | |
| 24 | Chimpanzee_Q14B(-) | 0.133 | 0.005 | 0.005 | 0.005 | 0.005 | 0.005 | 0.004 | 0.004 | 0.004 | 0.005 | 0.005 | 0.005 | 0.004 | 0.005 | 0.004 | 0.006 | 0.005 | 0.005 | 0.004 | 0.005 | 0.005 | 0.005 | 0.133 | 0.000 | | | | | | | | | | | | | | | | |
| 25 | Chimpanzee_Q14A(+) | 0.133 | 0.005 | 0.005 | 0.005 | 0.005 | 0.005 | 0.004 | 0.004 | 0.004 | 0.005 | 0.005 | 0.005 | 0.004 | 0.005 | 0.004 | 0.006 | 0.005 | 0.005 | 0.004 | 0.005 | 0.005 | 0.005 | 0.133 | 0.000 | 0.000 | | | | | | | | | | | | | | | |
| 26 | Chimpanzee_2_Not_in_palin | 0.003 | 0.132 | 0.132 | 0.132 | 0.131 | 0.132 | 0.131 | 0.132 | 0.131 | 0.132 | 0.132 | 0.132 | 0.132 | 0.133 | 0.132 | 0.133 | 0.132 | 0.132 | 0.131 | 0.132 | 0.133 | 0.133 | 0.003 | 0.132 | 0.132 | 0.000 | | | | | | | | | | | | | | |
| 27 | Chimpanzee_3_Not_in_palin | 0.132 | 0.007 | 0.006 | 0.006 | 0.006 | 0.006 | 0.006 | 0.006 | 0.006 | 0.006 | 0.007 | 0.007 | 0.006 | 0.007 | 0.006 | 0.007 | 0.007 | 0.007 | 0.006 | 0.006 | 0.007 | 0.007 | 0.132 | 0.006 | 0.006 | 0.131 | 0.000 | | | | | | | | | | | | | |
| 28 | Chimpanzee_Q10.9B(-) | 0.133 | 0.007 | 0.007 | 0.007 | 0.007 | 0.007 | 0.007 | 0.007 | 0.007 | 0.007 | 0.007 | 0.007 | 0.006 | 0.007 | 0.006 | 0.008 | 0.007 | 0.007 | 0.006 | 0.007 | 0.007 | 0.007 | 0.133 | 0.006 | 0.006 | 0.132 | 0.007 | 0.000 | | | | | | | | | | | | |
| 29 | Chimpanzee_Q10.9A(+) | 0.133 | 0.007 | 0.006 | 0.006 | 0.006 | 0.006 | 0.006 | 0.006 | 0.006 | 0.006 | 0.007 | 0.007 | 0.006 | 0.006 | 0.006 | 0.007 | 0.007 | 0.006 | 0.006 | 0.006 | 0.007 | 0.007 | 0.133 | 0.006 | 0.006 | 0.132 | 0.004 | 0.007 | 0.000 | | | | | | | | | | | |
| 30 | Chimpanzee_Q10.6B(-) | 0.133 | 0.006 | 0.006 | 0.006 | 0.006 | 0.006 | 0.006 | 0.006 | 0.006 | 0.006 | 0.006 | 0.006 | 0.006 | 0.006 | 0.006 | 0.007 | 0.006 | 0.006 | 0.006 | 0.006 | 0.007 | 0.007 | 0.133 | 0.005 | 0.005 | 0.132 | 0.004 | 0.007 | 0.000 | 0.000 | | | | | | | | | | |
| 31 | Chimpanzee_Q10.6A(+) | 0.133 | 0.007 | 0.007 | 0.007 | 0.007 | 0.007 | 0.007 | 0.007 | 0.007 | 0.007 | 0.007 | 0.007 | 0.006 | 0.007 | 0.006 | 0.008 | 0.007 | 0.007 | 0.006 | 0.007 | 0.007 | 0.007 | 0.133 | 0.006 | 0.006 | 0.132 | 0.007 | 0.000 | 0.007 | 0.007 | 0.000 | | | | | | | | | |
| 32 | Chimpanzee_Q10.Q14B(-) | 0.133 | 0.005 | 0.005 | 0.005 | 0.005 | 0.005 | 0.005 | 0.005 | 0.005 | 0.005 | 0.005 | 0.005 | 0.005 | 0.005 | 0.005 | 0.006 | 0.005 | 0.005 | 0.005 | 0.005 | 0.006 | 0.006 | 0.133 | 0.004 | 0.004 | 0.132 | 0.006 | 0.006 | 0.006 | 0.006 | 0.006 | 0.000 | | | | | | | | |
| 33 | Chimpanzee_Q10.Q14A(+) | 0.133 | 0.006 | 0.005 | 0.005 | 0.005 | 0.005 | 0.005 | 0.005 | 0.005 | 0.005 | 0.006 | 0.006 | 0.005 | 0.006 | 0.005 | 0.006 | 0.006 | 0.005 | 0.005 | 0.005 | 0.006 | 0.006 | 0.133 | 0.004 | 0.004 | 0.132 | 0.006 | 0.006 | 0.006 | 0.006 | 0.006 | 0.000 | 0.000 | | | | | | | |
| 34 | Chimpanzee_4_Not_in_palin | 0.003 | 0.132 | 0.132 | 0.132 | 0.132 | 0.132 | 0.132 | 0.132 | 0.132 | 0.132 | 0.133 | 0.133 | 0.132 | 0.133 | 0.132 | 0.133 | 0.133 | 0.132 | 0.132 | 0.133 | 0.133 | 0.133 | 0.003 | 0.132 | 0.132 | 0.001 | 0.132 | 0.132 | 0.132 | 0.133 | 0.132 | 0.132 | 0.132 | 0.000 | | | | | | |
| 35 | Chimpanzee_Q6.3B(-) | 0.133 | 0.007 | 0.007 | 0.007 | 0.007 | 0.007 | 0.007 | 0.007 | 0.007 | 0.007 | 0.007 | 0.007 | 0.007 | 0.007 | 0.007 | 0.008 | 0.007 | 0.007 | 0.007 | 0.007 | 0.007 | 0.007 | 0.134 | 0.006 | 0.006 | 0.132 | 0.007 | 0.002 | 0.007 | 0.006 | 0.002 | 0.006 | 0.006 | 0.133 | 0.000 | | | | | |
| 36 | Chimpanzee_Q6.3A(+) | 0.132 | 0.008 | 0.007 | 0.007 | 0.007 | 0.007 | 0.007 | 0.007 | 0.007 | 0.007 | 0.008 | 0.008 | 0.007 | 0.008 | 0.007 | 0.008 | 0.008 | 0.008 | 0.007 | 0.007 | 0.008 | 0.008 | 0.132 | 0.007 | 0.007 | 0.131 | 0.003 | 0.008 | 0.004 | 0.004 | 0.008 | 0.007 | 0.007 | 0.131 | 0.008 | 0.000 | | | | |
| 37 | Chimpanzee_Q4B(-) | 0.133 | 0.006 | 0.006 | 0.006 | 0.006 | 0.006 | 0.006 | 0.006 | 0.006 | 0.006 | 0.006 | 0.006 | 0.005 | 0.006 | 0.005 | 0.007 | 0.006 | 0.006 | 0.005 | 0.006 | 0.006 | 0.006 | 0.133 | 0.005 | 0.005 | 0.132 | 0.006 | 0.005 | 0.006 | 0.006 | 0.005 | 0.003 | 0.003 | 0.132 | 0.005 | 0.007 | 0.000 | | | |
| 38 | Chimpanzee_Q4A(+) | 0.133 | 0.006 | 0.005 | 0.005 | 0.005 | 0.005 | 0.005 | 0.005 | 0.005 | 0.005 | 0.006 | 0.006 | 0.005 | 0.006 | 0.005 | 0.006 | 0.006 | 0.005 | 0.005 | 0.005 | 0.006 | 0.006 | 0.133 | 0.002 | 0.002 | 0.132 | 0.006 | 0.005 | 0.005 | 0.005 | 0.005 | 0.004 | 0.005 | 0.132 | 0.006 | 0.007 | 0.004 | 0.000 | | |
| 39 | Chimpanzee_MtDNA | 0.542 | 0.504 | 0.504 | 0.504 | 0.504 | 0.504 | 0.503 | 0.503 | 0.503 | 0.503 | 0.505 | 0.505 | 0.504 | 0.505 | 0.504 | 0.504 | 0.504 | 0.504 | 0.503 | 0.505 | 0.505 | 0.505 | 0.542 | 0.505 | 0.505 | 0.542 | 0.506 | 0.505 | 0.506 | 0.506 | 0.505 | 0.505 | 0.506 | 0.543 | 0.505 | 0.507 | 0.506 | 0.506 | 0.000 | |
| 40 | Bonobo_MtDNA | 0.542 | 0.503 | 0.503 | 0.503 | 0.502 | 0.502 | 0.502 | 0.502 | 0.502 | 0.502 | 0.504 | 0.504 | 0.502 | 0.504 | 0.502 | 0.503 | 0.503 | 0.503 | 0.502 | 0.505 | 0.504 | 0.504 | 0.542 | 0.504 | 0.504 | 0.542 | 0.505 | 0.504 | 0.505 | 0.505 | 0.504 | 0.504 | 0.505 | 0.542 | 0.505 | 0.507 | 0.505 | 0.505 | 0.033 | 0.000 |

**Table S4. Palindromic NUMTs in chrY of bonobo and chimpanzee.** NUMTs are annotated with a corresponding palindrome arm (i.e., Q7 is the palindrome name and A indicates which arm). Sequence identity to mtDNA as well as the mtDNA coordinates with homology to the NUMT are included.

| Species | Chromosome | Start | End | NUMT length (bp) | Sequence identity to mtDNA (%) | MtDNA coordinates | Palindrome_annotation |
|---|---|---|---|---|---|---|---|
| Bonobo | chrY_pat_hsaY | 10798238 | 10802579 | 4341 | 74.155 | MT:16567-4348 | drome_but_similar_structure_2-numt-cluster |
| Bonobo | chrY_pat_hsaY | 10803193 | 10805524 | 2331 | 68.719 | MT:13702-16053 | drome_but_similar_structure_2-numt-cluster |
| Bonobo | chrY_pat_hsaY | 16903291 | 16907643 | 4348 | 75.496 | MT:1-4348 | Q20.9B_alone_3-numt-cluster |
| Bonobo | chrY_pat_hsaY | 16908259 | 16908488 | 229 | 70.815 | MT:15823-16054 | Q20.9B_alone_3-numt-cluster |
| Bonobo | chrY_pat_hsaY | 16908882 | 16910605 | 1723 | 69.954 | MT:13703-15438 | Q20.9B_alone_3-numt-cluster |
| Bonobo | chrY_pat_hsaY | 17600592 | 17602315 | 1723 | 70.011 | MT:13703-15438 | Q20.7B |
| Bonobo | chrY_pat_hsaY | 17602709 | 17602938 | 229 | 70.815 | MT:15823-16054 | Q20.7B |
| Bonobo | chrY_pat_hsaY | 17603554 | 17607907 | 4349 | 75.565 | MT:1-4348 | Q20.7B |
| Bonobo | chrY_pat_hsaY | 17708232 | 17712585 | 4349 | 75.542 | MT:1-4348 | Q20.7A |
| Bonobo | chrY_pat_hsaY | 17713201 | 17713429 | 228 | 70.386 | MT:15823-16054 | Q20.7A |
| Bonobo | chrY_pat_hsaY | 17713823 | 17715546 | 1723 | 70.011 | MT:13703-15438 | Q20.7A |
| Bonobo | chrY_pat_hsaY | 18336720 | 18338443 | 1723 | 69.977 | MT:13703-15438 | Q20.5B |
| Bonobo | chrY_pat_hsaY | 18338731 | 18339070 | 339 | 68.895 | MT:15714-16054 | Q20.5B |
| Bonobo | chrY_pat_hsaY | 18339686 | 18344039 | 4349 | 75.473 | MT:1-4348 | Q20.5B |
| Bonobo | chrY_pat_hsaY | 18444392 | 18448745 | 4349 | 75.473 | MT:1-4348 | Q20.5A |
| Bonobo | chrY_pat_hsaY | 18449361 | 18449700 | 339 | 68.895 | MT:15714-16054 | Q20.5A |
| Bonobo | chrY_pat_hsaY | 18449988 | 18451711 | 1723 | 69.977 | MT:13703-15438 | Q20.5A |
| Bonobo | chrY_pat_hsaY | 20627929 | 20629652 | 1723 | 69.977 | MT:13703-15438 | Q18.5B |
| Bonobo | chrY_pat_hsaY | 20629940 | 20630279 | 339 | 68.605 | MT:15714-16054 | Q18.5B |
| Bonobo | chrY_pat_hsaY | 20630895 | 20635248 | 4349 | 75.542 | MT:1-4348 | Q18.5B |
| Bonobo | chrY_pat_hsaY | 20735542 | 20739895 | 4349 | 75.519 | MT:1-4348 | Q18.5A |
| Bonobo | chrY_pat_hsaY | 20740511 | 20740850 | 339 | 68.895 | MT:15714-16054 | Q18.5A |
| Bonobo | chrY_pat_hsaY | 20741138 | 20742861 | 1723 | 69.977 | MT:13703-15438 | Q18.5A |
| Bonobo | chrY_pat_hsaY | 22935839 | 22937562 | 1723 | 69.977 | MT:13703-15438 | Q15.6B |

| | | | | | | | |
|---|---|---|---|---|---|---|---|
| Bonobo | chrY_pat_hsaY | 22937850 | 22938189 | 339 | 68.605 | MT:15714-16054 | Q15.6B |
| Bonobo | chrY_pat_hsaY | 22938805 | 22943158 | 4349 | 75.542 | MT:1-4348 | Q15.6B |
| Bonobo | chrY_pat_hsaY | 23043480 | 23047833 | 4349 | 75.496 | MT:1-4348 | Q15.6A |
| Bonobo | chrY_pat_hsaY | 23048449 | 23048788 | 339 | 68.605 | MT:15714-16054 | Q15.6A |
| Bonobo | chrY_pat_hsaY | 23049076 | 23050799 | 1723 | 69.977 | MT:13703-15438 | Q15.6A |
| Bonobo | chrY_pat_hsaY | 25545958 | 25547681 | 1723 | 69.862 | MT:13703-15438 | Q12.14B |
| Bonobo | chrY_pat_hsaY | 25547968 | 25548307 | 339 | 68.605 | MT:15714-16054 | Q12.14B |
| Bonobo | chrY_pat_hsaY | 25548923 | 25553276 | 4349 | 75.565 | MT:1-4348 | Q12.14B |
| Bonobo | chrY_pat_hsaY | 25653531 | 25657884 | 4349 | 75.565 | MT:1-4348 | Q12.14A |
| Bonobo | chrY_pat_hsaY | 25658500 | 25658839 | 339 | 68.605 | MT:15714-16054 | Q12.14A |
| Bonobo | chrY_pat_hsaY | 25659126 | 25660849 | 1723 | 69.862 | MT:13703-15438 | Q12.14A |
| Bonobo | chrY_pat_hsaY | 26297780 | 26299503 | 1723 | 69.977 | MT:13703-15438 | Q12.10B |
| Bonobo | chrY_pat_hsaY | 26299791 | 26300130 | 339 | 68.895 | MT:15714-16054 | Q12.10B |
| Bonobo | chrY_pat_hsaY | 26300746 | 26305098 | 4348 | 75.582 | MT:1-4348 | Q12.10B |
| Bonobo | chrY_pat_hsaY | 26405376 | 26409719 | 4339 | 75.468 | MT:1-4348 | Q12.10A |
| Bonobo | chrY_pat_hsaY | 26410335 | 26410674 | 339 | 68.605 | MT:15714-16054 | Q12.10A |
| Bonobo | chrY_pat_hsaY | 26410961 | 26412684 | 1723 | 69.862 | MT:13703-15438 | Q12.10A |
| Bonobo | chrY_pat_hsaY | 27029489 | 27031212 | 1723 | 69.977 | MT:13703-15438 | Q12.2B |
| Bonobo | chrY_pat_hsaY | 27031500 | 27031839 | 339 | 68.895 | MT:15714-16054 | Q12.2B |
| Bonobo | chrY_pat_hsaY | 27032455 | 27036809 | 4350 | 75.525 | MT:1-4348 | Q12.2B |
| Bonobo | chrY_pat_hsaY | 27137163 | 27141506 | 4339 | 75.422 | MT:1-4348 | Q12.2A |
| Bonobo | chrY_pat_hsaY | 27142122 | 27142461 | 339 | 68.497 | MT:15714-16054 | Q12.2A |
| Bonobo | chrY_pat_hsaY | 27142748 | 27144471 | 1723 | 69.862 | MT:13703-15438 | Q12.2A |
| Bonobo | chrY_pat_hsaY | 29532420 | 29534143 | 1723 | 69.954 | MT:13703-15438 | Q9.3B |
| Bonobo | chrY_pat_hsaY | 29534537 | 29534766 | 229 | 70.815 | MT:15823-16054 | Q9.3B |
| Bonobo | chrY_pat_hsaY | 29535382 | 29539735 | 4349 | 75.496 | MT:1-4348 | Q9.3B |
| Bonobo | chrY_pat_hsaY | 29640047 | 29644399 | 4348 | 75.513 | MT:1-4348 | Q9.3A |
| Bonobo | chrY_pat_hsaY | 29645015 | 29645244 | 229 | 70.815 | MT:15823-16054 | Q9.3A |
| Bonobo | chrY_pat_hsaY | 29645638 | 29647361 | 1723 | 70.011 | MT:13703-15438 | Q9.3A |
| Bonobo | chrY_pat_hsaY | 32069681 | 32071404 | 1723 | 69.977 | MT:13703-15438 | Q7B |

| | | | | | | | |
|---|---|---|---|---|---|---|---|
| Bonobo | chrY_pat_hsaY | 32071692 | 32072031 | 339 | 68.605 | MT:15714-16054 | Q7B |
| Bonobo | chrY_pat_hsaY | 32072647 | 32077002 | 4351 | 75.485 | MT:1-4348 | Q7B |
| Bonobo | chrY_pat_hsaY | 32170226 | 32174578 | 4348 | 75.536 | MT:1-4348 | Q7A |
| Bonobo | chrY_pat_hsaY | 32175194 | 32175533 | 339 | 68.605 | MT:15714-16054 | Q7A |
| Bonobo | chrY_pat_hsaY | 32175820 | 32177543 | 1723 | 69.805 | MT:13703-15438 | Q7A |
| Bonobo | chrY_pat_hsaY | 34586696 | 34588419 | 1723 | 69.862 | MT:13703-15438 | Q5B |
| Bonobo | chrY_pat_hsaY | 34588706 | 34589045 | 339 | 68.786 | MT:15714-16054 | Q5B |
| Bonobo | chrY_pat_hsaY | 34589661 | 34594004 | 4339 | 75.445 | MT:1-4348 | Q5B |
| Bonobo | chrY_pat_hsaY | 34694329 | 34698672 | 4339 | 75.445 | MT:1-4348 | Q5A |
| Bonobo | chrY_pat_hsaY | 34699288 | 34699627 | 339 | 68.786 | MT:15714-16054 | Q5A |
| Bonobo | chrY_pat_hsaY | 34699914 | 34701637 | 1723 | 69.862 | MT:13703-15438 | Q5A |
| Chimpanzee | chrY_hap2_hsaY | 5686494 | 5690835 | 4341 | 74.319 | MT:16555-4289 | Not_in_palindrome_3-numt-cluster |
| Chimpanzee | chrY_hap2_hsaY | 5691446 | 5691773 | 327 | 71.856 | MT:15666-15992 | Not_in_palindrome_3-numt-cluster |
| Chimpanzee | chrY_hap2_hsaY | 5692070 | 5693778 | 1708 | 70.213 | MT:13643-15375 | Not_in_palindrome_3-numt-cluster |
| Chimpanzee | chrY_hap2_hsaY | 6439768 | 6441350 | 1582 | 70.482 | MT:13785-15379 | Q14B |
| Chimpanzee | chrY_hap2_hsaY | 6441747 | 6441976 | 229 | 70.815 | MT:15763-15993 | Q14B |
| Chimpanzee | chrY_hap2_hsaY | 6442592 | 6446946 | 4350 | 75.507 | MT:1-4349 | Q14B |
| Chimpanzee | chrY_hap2_hsaY | 6547317 | 6551671 | 4350 | 75.507 | MT:1-4349 | Q14A |
| Chimpanzee | chrY_hap2_hsaY | 6552287 | 6552516 | 229 | 70.815 | MT:15763-15993 | Q14A |
| Chimpanzee | chrY_hap2_hsaY | 6552913 | 6554495 | 1582 | 70.482 | MT:13785-15379 | Q14A |
| Chimpanzee | chrY_hap2_hsaY | 11174210 | 11178549 | 4339 | 74.398 | MT:16555-4289 | t_in_palindrome_but_similar_3-numt-cluster |
| Chimpanzee | chrY_hap2_hsaY | 11179160 | 11179487 | 327 | 71.557 | MT:15666-15992 | t_in_palindrome_but_similar_3-numt-cluster |
| Chimpanzee | chrY_hap2_hsaY | 11179784 | 11181470 | 1686 | 69.217 | MT:13643-15375 | t_in_palindrome_but_similar_3-numt-cluster |
| Chimpanzee | chrY_hap2_hsaY | 11927477 | 11929059 | 1582 | 70.669 | MT:13785-15379 | drome_but_similar_structure_3-numt-cluster |
| Chimpanzee | chrY_hap2_hsaY | 11929456 | 11929685 | 229 | 70.815 | MT:15763-15993 | drome_but_similar_structure_3-numt-cluster |
| Chimpanzee | chrY_hap2_hsaY | 11930301 | 11934652 | 4347 | 75.684 | MT:1-4349 | drome_but_similar_structure_3-numt-cluster |
| Chimpanzee | chrY_hap2_hsaY | 15509776 | 15511344 | 1568 | 69.981 | MT:13785-15379 | Q10.9B |
| Chimpanzee | chrY_hap2_hsaY | 15511741 | 15511970 | 229 | 70.815 | MT:15763-15993 | Q10.9B |
| Chimpanzee | chrY_hap2_hsaY | 15512586 | 15516918 | 4328 | 75.572 | MT:1-4327 | Q10.9B |
| Chimpanzee | chrY_hap2_hsaY | 15617350 | 15621702 | 4348 | 75.524 | MT:1-4349 | Q10.9A |

| | | | | | | | |
|---|---|---|---|---|---|---|---|
| Chimpanzee | chrY_hap2_hsaY | 15622318 | 15622547 | 229 | 70.815 | MT:15763-15993 | Q10.9A |
| Chimpanzee | chrY_hap2_hsaY | 15622944 | 15624526 | 1582 | 70.482 | MT:13785-15379 | Q10.9A |
| Chimpanzee | chrY_hap2_hsaY | 16246574 | 16248156 | 1582 | 70.482 | MT:13785-15379 | Q10.6B |
| Chimpanzee | chrY_hap2_hsaY | 16248553 | 16248782 | 229 | 70.815 | MT:15763-15993 | Q10.6B |
| Chimpanzee | chrY_hap2_hsaY | 16249398 | 16253750 | 4348 | 75.524 | MT:1-4349 | Q10.6B |
| Chimpanzee | chrY_hap2_hsaY | 16354148 | 16358480 | 4328 | 75.572 | MT:1-4327 | Q10.6A |
| Chimpanzee | chrY_hap2_hsaY | 16359096 | 16359325 | 229 | 70.815 | MT:15763-15993 | Q10.6A |
| Chimpanzee | chrY_hap2_hsaY | 16359722 | 16361290 | 1568 | 69.981 | MT:13785-15379 | Q10.6A |
| Chimpanzee | chrY_hap2_hsaY | 16981483 | 16983051 | 1568 | 70.044 | MT:13785-15379 | Q10.1B |
| Chimpanzee | chrY_hap2_hsaY | 16984293 | 16988622 | 4325 | 75.596 | MT:1-4327 | Q10.1B |
| Chimpanzee | chrY_hap2_hsaY | 17085779 | 17090105 | 4322 | 75.55 | MT:1-4327 | Q10.1A |
| Chimpanzee | chrY_hap2_hsaY | 17091347 | 17092915 | 1568 | 70.044 | MT:13785-15379 | Q10.1A |
| Chimpanzee | chrY_hap2_hsaY | 20640274 | 20644614 | 4340 | 74.358 | MT:16555-4289 | drome_but_similar_structure_3-numt-cluster |
| Chimpanzee | chrY_hap2_hsaY | 20645225 | 20645552 | 327 | 71.557 | MT:15666-15992 | drome_but_similar_structure_3-numt-cluster |
| Chimpanzee | chrY_hap2_hsaY | 20645849 | 20647557 | 1708 | 70.196 | MT:13643-15375 | drome_but_similar_structure_3-numt-cluster |
| Chimpanzee | chrY_hap2_hsaY | 21394319 | 21395901 | 1582 | 70.482 | MT:13785-15379 | Q6.3B |
| Chimpanzee | chrY_hap2_hsaY | 21397143 | 21401475 | 4328 | 75.647 | MT:1-4327 | Q6.3B |
| Chimpanzee | chrY_hap2_hsaY | 21501796 | 21506146 | 4346 | 75.684 | MT:1-4349 | Q6.3A |
| Chimpanzee | chrY_hap2_hsaY | 21506762 | 21506991 | 229 | 70.815 | MT:15763-15993 | Q6.3A |
| Chimpanzee | chrY_hap2_hsaY | 21507388 | 21508970 | 1582 | 70.607 | MT:13785-15379 | Q6.3A |
| Chimpanzee | chrY_hap2_hsaY | 23845108 | 23846676 | 1568 | 70.106 | MT:13785-15379 | Q4B |
| Chimpanzee | chrY_hap2_hsaY | 23847918 | 23852248 | 4326 | 75.687 | MT:1-4327 | Q4B |
| Chimpanzee | chrY_hap2_hsaY | 23952662 | 23957016 | 4350 | 75.53 | MT:1-4349 | Q4A |
| Chimpanzee | chrY_hap2_hsaY | 23958258 | 23959816 | 1558 | 70.006 | MT:13797-15379 | Q4A |
